# Recombinogenic G-quadruplexes in the Newtonian DNA Sequence Space

**DOI:** 10.64898/2026.06.30.735570

**Authors:** Vitaly Kuryavyi

## Abstract

The universe of possible nucleotide sequences expands combinatorially with sequence length, vastly exceeding the fraction sampled by real genomes. Yet genomic sequences exhibit reproducible compositional symmetries and recurrent structural motifs, indicating that biological sequence space is shaped by strong organizing constraints. Here, we introduce an explicit framework for constructing and visualizing the complete sequence universe using the Newtonian polynomial for a four-letter alphabet, and for identifying biologically relevant subsets through the application of fundamental filters.

Three filters of biological relevance are formulated: (i) the constraint that DNA predominantly exists as an antiparallel-stranded double helix, (ii) the second Chargaff parity rule, which enforces approximate strand symmetry in single-stranded sequence composition, and (iii) genome shadows, reflecting the imprint of concerted sequence changes. Successive application of these filters dramatically reduces the accessible sequence space and reveals distinct symmetry classes.

Among these, mirror-symmetric sequences occupy a privileged position because they are invariant under strand reversal and therefore compatible with both antiparallel and parallel strand orientations. This dual compatibility enables such sequences to bridge otherwise disjoint structural subspaces of DNA. G-rich members of this class are shown to have a strong propensity to form G-quadruplex architectures that incorporate parallel-stranded domains while remaining compatible with duplex DNA. We propose that this structural versatility provides a mechanistic basis for the recurrent association of G-rich mirror-symmetric sequences with recombination hotspots and genome rearrangements. Together, these results establish a symmetry-based framework for understanding how combinatorial sequence space is filtered into biologically functional DNA motifs.

## Inevitability of pattern formation in sequence space

In processes governing the flow of genetic information, DNA and RNA explore sequence space through three principal mechanisms: (i) changes in genome size, (ii) local sequence alteration by mutation, and (iii) recombination, which reorders extended fragments within or between genomes. The theoretical space of nucleotide sequences is infinite and countable, yet it is constructed from a markedly reduced four-letter alphabet. As a consequence, in sufficiently long nucleotide strings functioning in biological systems, the emergence of simple, repetitive, and symmetric patterns is not exceptional but inevitable. This inevitability follows directly from combinatorial constraints and does not rely on assumptions about selective advantage or functional annotation.

## Repetitive DNA as a combinatorial consequence, not an anomaly

Early molecular biology, largely focused on nucleic acids as carriers of vertically and horizontally transmitted information, treated repetitive DNA sequences in viral and bacterial genomes as peripheral despite their substantial genomic presence. In mammalian genomes, where repetitive sequences constitute a dominant fraction, their prevalence proved harder to ignore and led to the introduction of terms such as “selfish” or “junk” DNA(1,2), implying copying-driven accumulation as their primary explanation. While cellular molecular machinery is indeed highly optimized for copying, the more fundamental question is whether genomes—very long texts written with only four symbols—can in principle be free of repetitive features. Stated differently, is it possible for such texts to consist entirely of unique sequence patterns, or are repetitions and symmetries unavoidable fossils of the finite alphabet itself? Addressing this question requires stepping outside individual genomes and considering the full space of theoretically possible sequences.

## Limitations of genome-centric analyses and motivation for a global view

The full sequence space is commonly invoked either as a reservoir from which evolution samples genetic material or as a mathematical background for formulating statistical hypotheses tested on real genomes. Numerous analytical frameworks—such as Z-curves(3), multidimensional walks(4), and related compositional analyses—have revealed striking symmetries and constraints in nucleotide composition and ordering, including those associated with the second Chargaff parity rule(5–11). However, these approaches typically retain a dual focus on real genomes versus randomized controls, leaving the remaining, overwhelmingly vast universe of possible sequences structurally unexamined. In this work, I propose an explicit visualization and organization of the complete sequence space, structured in a way that places real genomes within its full combinatorial context rather than treating them in isolation.

## Instrument: explicit construction of the full sequence space

To address these limitations, I introduce a framework for explicitly constructing and visualizing the complete universe of nucleotide sequences. This universe can be represented mathematically using the Newtonian polynomial (equivalently, the multinomial expansion for four variables), which enumerates all possible sequences of a given length composed of the four nucleotides. When plotted graphically, for example as pyramidal or higher-dimensional lattice representations, this construction makes the intrinsic symmetries of sequence space directly visible. Importantly, this representation is agnostic to biological function: it treats all sequences as equal members of the combinatorial universe. Real genomic sequences, when mapped onto this space, occupy an exceedingly small and highly structured subset, revealing that their properties are not arbitrary but constrained by deeper organizing principles.

## Filters of biological relevance and emergence of symmetry classes

Biological sequences do not sample this universe freely. Instead, they are constrained by a small number of fundamental rules that act as filters, progressively restricting the accessible subspace. In this work, three such rules are formulated explicitly: (i) DNA exists predominantly as an antiparallel-stranded double helix, imposing orientation-dependent complementarity constraints; (ii) the second Chargaff parity rule, which enforces approximate strand symmetry in single-stranded sequence composition; and (iii) genome shadows, reflecting the imprint of concerted changes imposed on functionally active sequences. When these filters are applied to the full sequence space, distinct symmetry classes emerge naturally. Among them, sequences with mirror symmetry occupy a privileged position, as they can bridge subspaces defined by antiparallel and parallel strand orientations. These sequences are therefore predisposed to adopt alternative DNA architectures and to participate in genome rearrangement processes.

## Structural realizability of symmetry classes

The symmetry classes that emerge from the filtered sequence space are not merely abstract mathematical constructs but correspond to distinct sets of physically realizable DNA architectures. The canonical antiparallel double helix occupies only one such subspace. Parallel strand orientations, although energetically disfavored in standard duplex DNA, are well established in non-canonical nucleic acid structures, including triplexes and quadruplexes. Sequences possessing mirror symmetry are unique in this regard: because their sequence identity is preserved under reversal, they are compatible with both antiparallel and parallel strand alignments. This dual compatibility allows mirror-symmetric sequences to function as structural junctions between otherwise disjoint regions of sequence space, expanding the repertoire of accessible DNA topologies under physiological conditions.

## G-rich sequences as recombinogenic hubs

Among mirror-symmetric sequences, G-rich motifs are of particular interest because of their propensity to form G-quadruplex structures that naturally incorporate parallel-stranded domains. These structures are polymorphic, with loop lengths and strand orientations that can vary while preserving a common G-tetrad core. As a result, G-rich mirror-symmetric sequences are capable of transitioning between duplex and quadruplex states and of accommodating both parallel and antiparallel DNA elements within a single structural framework. This architectural flexibility suggests a mechanistic basis for their recurrent association with genome rearrangements and recombination hotspots: rather than serving as passive repetitive elements, such sequences may act as structural hubs that transiently stabilize alternative DNA conformations required for strand exchange and genome reorganization(12,13).

## Results and Discussion

### 1. Genomes as nucleotide sequences or sets of nucleotide sequences

The sequence of a single strand of any chromosome—whether already sequenced or yet to be determined—corresponds to one element within the complete universe of nucleotide sequences of a given length. This universe is exhaustively enumerated by the Newtonian polynomial (multinomial expansion) for a four-letter alphabet,

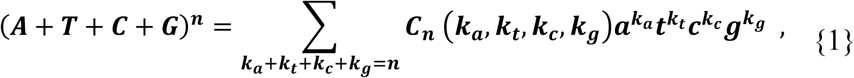

where (n) denotes sequence length and the multinomial coefficient is defined as

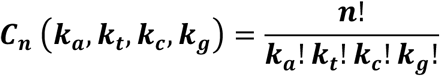

Each term of this expansion represents a class of sequences sharing identical nucleotide composition, while individual sequences within a class differ by the order of bases along the strand.

For short sequence lengths, the complete sequence space can be written explicitly. For dinucleotides ((n=2)),

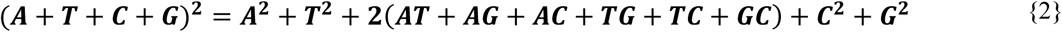

and for trinucleotides ((n=3)),

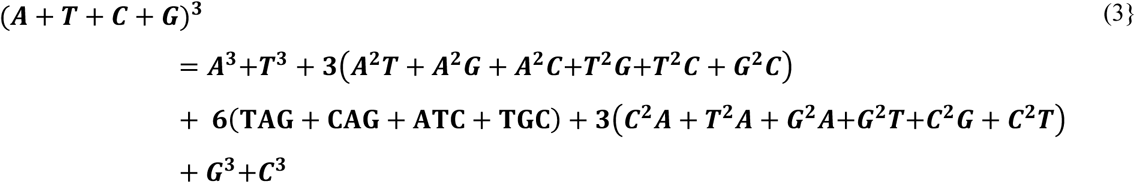

These expansions are represented graphically in Figure 1 as pyramidal lattices, in which vertices correspond to nucleotides and edges represent their sequential ordering along a strand. The apex of each pyramid has four connections and is therefore used to denote the 5′ end of a sequence, while the four vertices at the base correspond to possible 3′ termini. Separate pyramids with A, T, G, or C at the apex distinguish sequences with identical composition but different base order. For example, the dinucleotides 5′AT and 5′TA—members of the same compositional class (2(AT)) in Eq. {2}—occupy distinct positions in the A- and T-pyramids, respectively (Figure 1A).

**Figure 1.**
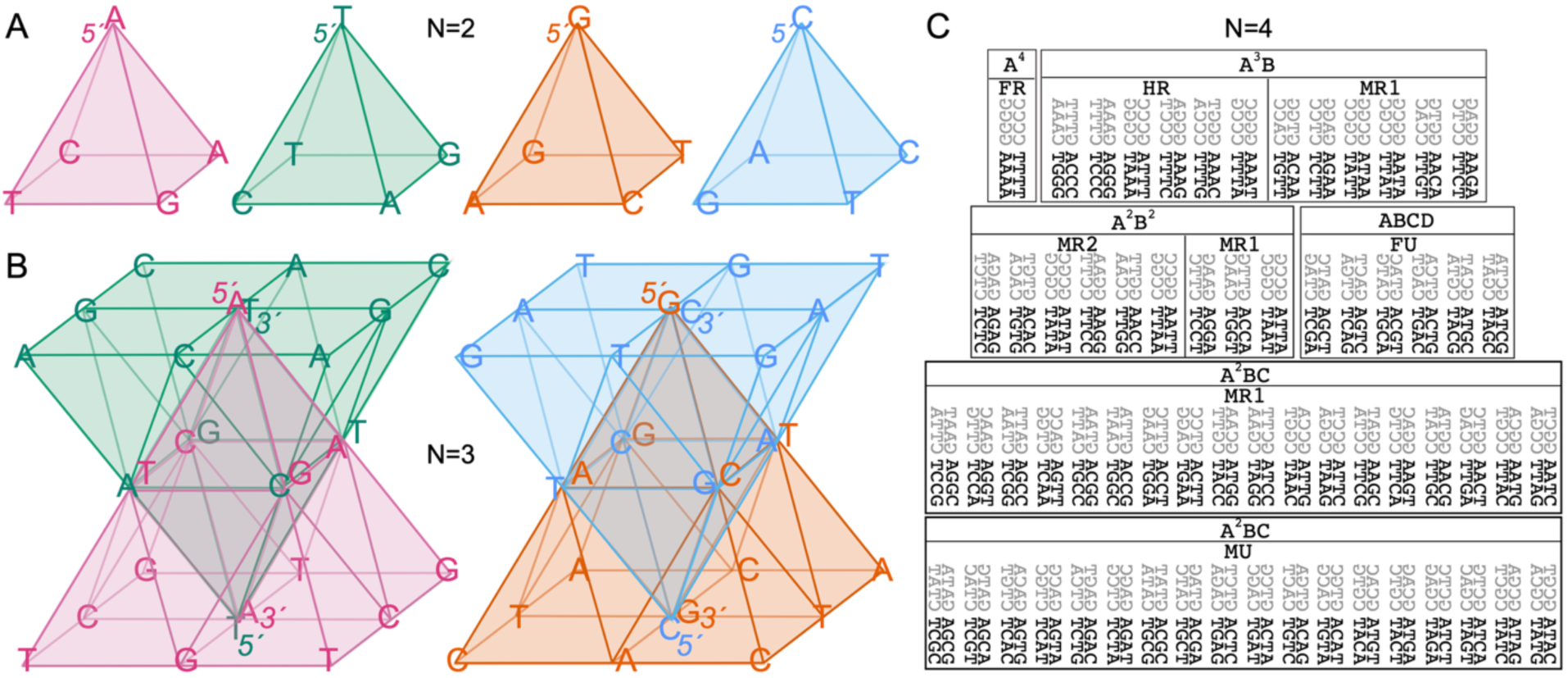
Full combinatorial sequence space of di-, tri-, and tetranucleotides. Di- and trinucleotides are presented as pyramids, whereas tetranucleotides are represented as complementary duplexes arranged in pairwise juxtapositions related by concerted A↔G and C↔T substitutions (“shadow” transformations). Tetranucleotides are further grouped according to sequence complexity (see text). **(A) Dinucleotide pyramids.** The sixteen dinucleotides are organized into four pyramids (A-, C-, G-, and T-pyramids). Solid lines connect the two nucleotides within each dinucleotide. The apical nucleotide defines both the pyramid type and the 5′ end of the sequence, which is read from the apex to each of the four nucleotides at the base. **(B) Trinucleotide pyramids.** Expansion of the dinucleotide pyramids generates paired trinucleotide pyramids. The A- and T-pyramids, as well as the G- and C-pyramids, form complementary pairs in which one pyramid is inverted relative to the other. **(C) Tetranucleotide sequence space.** Tetranucleotides are shown as complementary duplexes and paired with corresponding duplexes related by reflection about the horizontal axis, producing the shadow transformation generated by concerted A↔G and C↔T substitutions. The complete set is partitioned into polynomial classes (A₃B, A₂B₂, A₂BC, and ABCD) and further subdivided into the **Fully Repetitive (FR), Highly Repetitive (HR), Moderately Repetitive 1 (MR1), Moderately Repetitive 2 (MR2), Moderately Unique (MU),** and **Fully Unique (FU)** sequence classes described in the text.

For trinucleotides, the pyramids are rotated and extended (Figure 1B) to emphasize their ability to grow naturally into higher-order oligomers (4-mers, 5-mers, …, (n)-mers). Pairs of pyramids are further merged to compactly represent the full space of antiparallel-stranded DNA duplexes. In this representation, complementary sequences are read along the same edges of the merged structure in opposite orientations from apex to base within a single color. For example, the mirror-symmetric trinucleotide 5′AGA3′ is paired with its complement 5′TCT3′ within the central bipyramid (Figure 1B). The same complementary relation holds for the pairs of non-symmetric trinucleotide sequences outside the central bipyramid. This case requires transfer of corresponding trajectories between complementary bipyramids, indicating that the full antiparallel symmetry group is distributed across the paired structures rather than contained within a single geometric object. For example, AAG trinucleotide in the left bipyramid, has its complement as CTT in the bipyramid on the right. For any odd sequence length, the apex of one pyramid meets the center of the base of its complementary inverted pyramid, reflecting the inherent pairing of antiparallel strands. Two merged pyramid pairs thus represent the complete Newtonian sequence space for (n=3) constrained to antiparallel duplex DNA, initiating with either AT (in the bipyramid on the left) or GC (in the bipyramid on the right) base pairs.

In contrast to odd sequence lengths, for even sequence lengths, certain sequence trajectories become self-conjugate under antiparallel reversal, so that a sequence and its antiparallel complement occupy the same geometric edge. These self-conjugate antiparallel trajectories produce unavoidable edge degeneracy, requiring the doubling of specific edges in the graphical representation (Figure S0 showing inversion of dinucleotide pyramids). Thus, odd-length spaces exhibit vertex-centered trajectory pairing, whereas even-length spaces exhibit edge-centered symmetry with fixed trajectories under reversal. To avoid artificial doubling of the degenerate sequences in the polynomial for the even sequence lengths for the sake of satisfying antiparallel pairing, for the case of tetranucleotides, the full sequence space is presented in a tabular format (Figure 1C). For longer sequence lengths, the orbit formalism will be introduced later.

This spatial construction makes symmetry relationships within sequence space immediately apparent. For every position occupied by adenine in one pair of pyramids, guanine appears in a symmetrically related position in the other; an analogous correspondence holds for thymine and cytosine (Figure 1B). Examination of di- and trinucleotide patterns reveals distinct subclasses of sequences. These include fully repetitive sequences composed of a single nucleotide (e.g., A^2^, G^2^), fully unique sequences, and intermediate classes in which some—but not all—letters are, repeated. In trinucleotides, such moderately repetitive subclasses emerge naturally, for example in terms such as 3(A^2^T+A^2^G+A^2^C+T^2^G+T^2^C+G^2^C) in Eq. (14). For tetranucleotides and longer sequences, these distinctions expand into recognizable groups of fully repetitive (FR), highly repetitive (HR), moderately repetitive (MR1 and MR2), moderately unique (MU), and fully unique (FU) sequences (Figure 1C).

### 1.2 Replication and recombination in the full combinatorial space

Recombination between carriers of nucleotide sequences is commonly perceived as a mechanism that generates diversity, while replication—subject to mutation—is viewed as the process by which selected, functional genomes are propagated across generations. These views are appropriate when individual evolutionary events are considered in isolation. However, when replication and recombination are examined within the context of the full combinatorial sequence space, their semantic meaning changes in a fundamental way.

To illustrate this point, consider the complete set of trinucleotides, Eq (14), treated hypothetically as a population of single-stranded “genomes,” with individual nucleotides acting as the simplest possible “genes.” In this population of 64 sequences, more than half (37 out of 64) contain at least one copy of the “gene”—A, C, G, or T. Importantly, the presence of the “genes” in these “genomes” does not result from copying or inheritance from another sequence but arises purely as a consequence of the combinatorial structure of sequence space itself. This example illustrates a distinct fundamental feature of the sequence space: the diversity.

A second thought experiment further emphasizes the distinction between sequence composition and population diversity. Consider two trinucleotide genomes, AA|T and TT|A, undergoing a recombination event at the indicated junction. Recombination produces the sequences AAA and TTT. This outcome is indistinguishable from the result of error-free replication of genomes AAA and TTT. At the level of individual sequences, recombination has produced valid members of the sequence space. At the population level, however, diversity has decreased: the sequences AAT and TTA are lost, while AAA and TTT now occur with doubled frequency. Thus, recombination does not necessarily increase diversity; instead, it redistributes occupancy within sequence space. This thought experiment highlights that “sequence” and “diversity” are distinct yet inseparable properties of populations embedded in finite combinatorial spaces.

Having established these principles in a simplified setting, the natural question is whether real genomes can approach the limits of sequence space occupancy. For example, the chromosome of *Escherichia coli* K-12 has a length of approximately 4,639,676 nucleotides. The corresponding Newtonian sequence space, (a+t+c+g)^4,639,676^, is astronomically large, but the number of sequences representing a single, mutation-free circular genome is itself nontrivial. A circular genome of this length corresponds to 4,639,676 distinct linear single-stranded sequences, and together with the same number of complementary sequences, yields 9,279,352 sequence representations for a single genomic composition class. Even this number must be subtracted from the vastly larger total number of sequences encompassed by the full polynomial expansion.

Despite this immense combinatorial freedom, it is immediately evident that many sequences of length 4,639,676 are incompatible with biological function. Trivial examples include homopolymeric sequences composed entirely of A, C, G, or T, as well as near-homopolymers interrupted by only one or a few nucleotides. Such sequences can be excluded by elementary considerations of chemical stability and information content. However, as sequence complexity increases, these intuitive filters rapidly lose discriminative power. This raises a more fundamental question: can biologically compatible sequences be distinguished from non-compatible ones by general rules, rather than by case-specific functional annotation? In other words, are there intrinsic properties of DNA sequences derived from living organisms that act as global filters, carving structured voids within the densely packed combinatorial sequence space?

### 2.1. Results. Filters

#### 2.1 Antiparallel-stranded duplex DNA with A–T and G–C base pairs

At first glance, it may not appear obvious that the antiparallel double helix itself functions as a *filter* on sequence space. Although the canonical DNA double helix(15) does not impose a specific sequence motif requirement for adopting the structure, it does impose strong constraints on the repertoire of sequences compatible with biological function.

To illustrate this point, it is instructive to consider a contrasting alternative: **parallel-stranded double-stranded DNA (ps-DNA)**. From a structural perspective, ps-DNA can be constructed using either *trans* or *cis* base pairs (16–19). In a trans A–T base pair (Figure 2A, left), the sugar residues of the paired nucleotides are co-oriented. In contrast, in a cis Watson–Crick base pair, one nucleoside must adopt a syn glycosidic conformation to achieve sugar co-orientation, as observed in Z-DNA (17,20)

**Figure 2.**
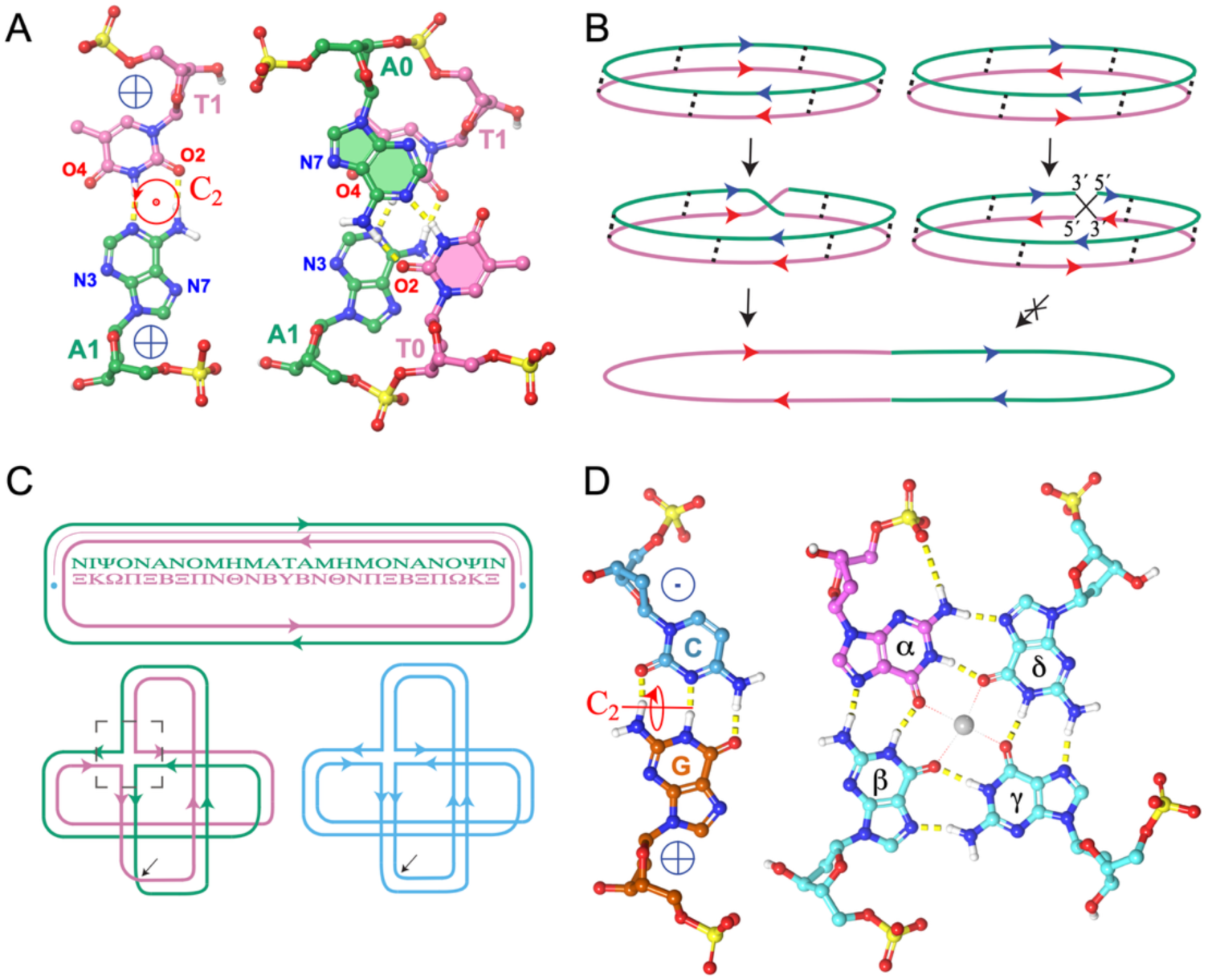
Parallel-stranded DNA, but not antiparallel-stranded DNA, can undergo topological transformations leading to self-complementary single-stranded structures with Möbius-strip topology and provides a conceptual bridge to G-quadruplex architecture. (A) Pseudo-symmetry of trans base pairs. Trans A·T and G·C base pairs possess approximate C₂ symmetry with the symmetry axis perpendicular to the base-pair plane (left). Sugar directions are indicated by symbols denoting the orientation of the 5′→3′ backbone relative to the viewer. Two trans A·T base pairs can be connected to form a parallel-stranded duplex in which one base pair is rotated by 180°, generating the parallel-stranded duplex 5′-T_0_A_1_-3′/5′-A_0_*T*_1_-3′ (right). All nucleotides shown in panels A–C adopt anti glycosidic conformations. **(B) Topological conversion of parallel-stranded DNA.** A parallel-stranded duplex (left), but not an antiparallel-stranded duplex (right), permits strand cleavage, 180° rotation, and religation. This transformation produces a single-stranded circular molecule of doubled length in which one half of the sequence is complementary to the other in the parallel orientation, generating a Möbius-strip topology. **(C) Mirror-symmetric circular DNA and strand-crossing transformation.** A circular antiparallel duplex containing a mirror-symmetric sequence is shown in the upper panel. Conversion into a junction between parallel-stranded and antiparallel-stranded duplexes is illustrated in the lower left panel; the two original strands are colored green and magenta. The black arrow marks the position at which the parallel-stranded segment can be cleaved and rotated by 180°. Religating the strands produces a single continuous strand of doubled length (blue) with Möbius-strip topology. **(D) Relationship between cis base pairing and G-tetrads.** A cis G·C base pair exhibits approximate C₂ symmetry about an axis lying within the base-pair plane (left). A G-tetrad is shown on the right, with α-guanine (magenta) adopting the syn glycosidic conformation, whereas β-, γ-, and δ-guanines adopt anti conformations. The α/β guanine pair is isomorphic to a C·G base pair, with the syn conformation reversing the relative sugar orientation. This structural correspondence provides a geometric connection between Watson–Crick duplexes and G-quadruplex architecture. **In panels A and D, 5′→3′ sugar directions are indicated by symbols representing backbone orientation relative to the viewer: ⊙ (toward the viewer) and ⊗ (away from the viewer).**

A key distinction between trans- and cis-base pairs lies in the orientation of the pseudo-symmetry axis relating the C–N glycosidic bonds of the two nucleosides. In trans base pairs, this axis is perpendicular to the plane of the base pair (Figure 2A), whereas in cis base pairs it lies within the plane of the base pair (Figure 2D, left). In ps-DNA constructed from trans base pairs, this symmetry leads to equivalence of the two grooves. For example, the N3/O4 groove of base pair A₁T₁ continues smoothly into the opposing O2/N7 groove of the neighboring base pair A₀T₀ (Figure 2A), with the two base pairs related by a 180° rotation. In the C·C⁺ base pairs of the i-motif, the two grooves are indistinguishable because the base pair itself is symmetric (21).

These two structural features—rotational symmetry and co-oriented sugar direction—lead to a striking topological consequence. Two strands of such parallel-stranded DNA can be broken at any single point, rotated by 180°, and reconnected while preserving the overall structure. Upon reconnection, however, the two strands become topologically fused into a single strand with **Möbius strip topology** (Figure 2B, left). In contrast, for antiparallel-stranded DNA, a 180° rotation would juxtapose 3′–3′ and 5′–5′ termini, a configuration incompatible with natural covalent linkage. Reconnection of antiparallel strands therefore requires a full 360° rotation (Figure 2B, right).

It follows that parallel-stranded DNA possesses an intrinsic propensity for strand crossover and self-recombination, facilitating rearrangement and shuffling of genetic material, as illustrated for the circular case in Figure 2B. After a single such event, the length of the resulting single strand becomes doubled. One half of this strand is now parallel-complementary to the other half. A minimal illustrative example is the tetranucleotide 5′-AGTC-3′, which can be divided into two halves, 5′-AG-3′ and 5′-TC-3′, related by parallel complementarity. Consistent with this reasoning, A–T and G–C trans base pairs are known to support the formation of parallel-stranded duplex DNA (22).

From a combinatorial perspective, every single strand among (a+t+c+g)^4,639,676^ possible variants can be represented as two contiguous halves joined into continuity. This yields (a+t+c+g)^2,319,838^ theoretically possible half-sequences. For each such half, there exists exactly one parallel-complementary partner. Consequently, out of the full (a+t+c+g)^4,639,676^ sequence space, (a+t+c+g)^2,319,838^ sequences constitute candidates to be excluded by **Rule 1**, the requirement for antiparallel-stranded DNA.

Genomes in which one half of the strand is parallel-complementary to the other half are biologically incompatible, because all known replicative DNA systems rely on antiparallel duplexes with A–T and G–C complementarity. This thought experiment further suggests that, relative to antiparallel-stranded DNA, parallel-stranded DNA is less suited for long-term preservation of sequence information and is instead predisposed toward sequence diversification and recombination.

### 2.2 The Second Chargaff Parity Rule

The first Chargaff rule established an approximate balance between A/T and G/C base contents in double-stranded DNA(23). The **second Chargaff parity rule (SCPR)** extends this observation to single DNA strands, stating that, to a close approximation, A ≈ T and G ≈ C within each strand (10). This rule holds with remarkable precision in bacterial genomes(9).

Two operational forms of SCPR have been distinguished: one applying to **mononucleotides (C^II^mono)** and the other to **oligonucleotides (C^II^oligo)**(5). Both forms have been examined extensively in bacterial and eukaryotic genomes and were confirmed for oligonucleotide lengths ranging from 3 to 9 nucleotides(7,24) (6,8). According to **the C^II^oligo** rule, single strands of double-stranded genomes are composed of pairwise equal numbers of **antiparallel-complementary k-mers**.

The effect of applying the second Chargaff parity rule as a *filter* can be illustrated explicitly using the complete sequence space at length four. All constituent ((A+T+C+G)^4^ =256) oligonucleotides are shown in Figure 1C as A/T- and G/C-paired duplexes arranged in a symmetric juxtaposition. Equivalent base positions in black and gray duplex pairs are related by C↔T and A↔G substitutions reflected about the horizontal axis separating these two pairs of duplexes. For illustrative purposes only, each duplex in this set is referred to here as a “genome.”

This complete group can be subdivided based on sequence composition and internal repetition patterns. Specifically, it contains 2 duplex pairs in the A^₄^ group, 12 pairs in the A^₃^B group, 18 pairs in the A^₂^B^₂^ group, 36 pairs in the A^₂^BC group, and 6 pairs in the ABCD group.

The A^₄^ group consists of **Fully Repetitive (FR)** sequences, composed entirely of a single nucleotide. One half of the A^₃^B group is classified as **Highly Repetitive (HR)**, containing three identical bases in a row and a different base at one end. The other half belongs to **Moderately Repetitive Group 1 (MR1)**, distinguished by the placement of two identical adjacent bases, either at one or both ends or in the middle of the sequence. By the same criterion, one-third of the A^₂^B^₂^ group also belongs to MR1.

The remaining two-thirds of the A^₂^B^₂^ group form **Moderately Repetitive Group 2 (MR2)**. MR2 sequences are composed of two repetitive halves, which may be identical or different (e.g., ACAC or AAGG). MR1-type sequences also account for half of the A₂BC group. The remaining half of A^₂^BC comprises **Moderately Unique (MU)** sequences, which contain at least one repeated base, but in which identical bases are always separated along the sequence. Finally, the ABCD group consists of six **Fully Unique (FU)** oligonucleotide duplex pairs, along with their six symmetric reflections.

Application of the **C^II^mono filter** to the full set eliminates all sequences except those belonging to the FU (ABCD) group, as only in this group does the condition A = T and G = C hold within single strands. Subsequent application of **Filter 1** (the requirement for antiparallel-stranded DNA) further eliminates sequences such as ACTG/TGAC, GTCA/CAGT, AGTC/TCAG, and GACT/CTGA, because these are composed of two halves related by *parallel* complementarity. As a result, only **four duplex pairs**, corresponding to **16 individual sequences**, pass both filters for biological compatibility out of the original 256-member sequence space.

The purpose of the tetranucleotide table is not exhaustive classification per se, but to provide the shortest sequence length at which the dramatic contraction of sequence space under biologically relevant filters becomes directly visible.

The number of sequences satisfying the C^II^mono condition can be obtained directly from the multinomial expansion. For a sequence of length *k*, the constraint imposed by the rule requires:

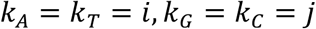

with

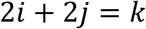

Therefore:

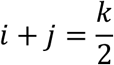

No solutions exist for odd k, indicating that exact C^II^mono symmetry is only possible for even-length sequences. Defining:

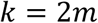

all valid sequence compositions are indexed by:

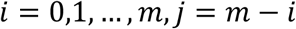

For each admissible pair (*i,j*), the number of sequences is given by the multinomial coefficient:

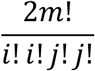

Summing over all allowed values yields the total number of C^II^mono-compliant sequences:

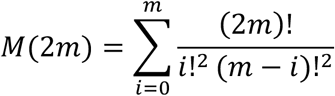

Although the absolute number of C^II^mono-compatible sequences increases rapidly with sequence length, their fraction within the full sequence universe:

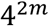

decreases sharply, demonstrating that the C^II^mono rule defines an increasingly sparse symmetry-constrained subspace within the Newtonian sequence universe.

Exact C^II^mono symmetry is only possible for even-length sequences. For odd lengths, parity constraints require a minimal imbalance between one pair of complementary nucleotides. The biologically relevant extension of the rule therefore corresponds to sequences in which one complementary pair differs by a single nucleotide while the second pair remains balanced. In this formulation, odd-length sequences represent the nearest parity-compatible realization of the C^II^mono constraint.

So, for odd lengths, the biologically meaningful generalization is:

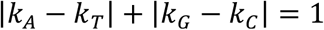

or equivalently:
one complementary pair differs by exactly 1, the other remains equal.

Let

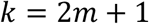

Then the imbalance must occur in exactly one complementary pair. So there are two possibilities:

Case 1

A/T carries the excess nucleotide:

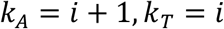

while:

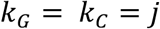

with:

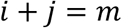

The number of sequences is:

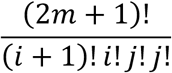

Case 2

G/C carries the excess nucleotide:

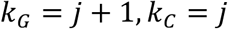

with:

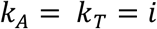

giving:

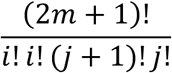

Because the two cases are symmetric, the total becomes:

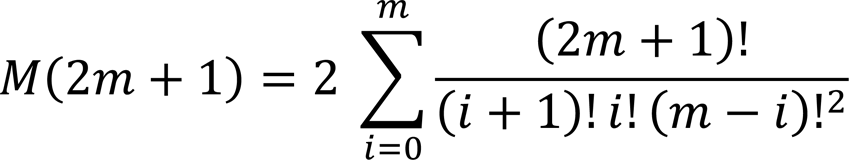

Insertions and deletions therefore represent more than local modifications of nucleotide composition. By changing sequence parity, they force transitions between exact and parity-limited realizations of the C^II^mono constraint, thereby transferring sequences between distinct symmetry classes of the filtered sequence universe. The distinction between the sequences with odd and even lengths has already been noted when constructing pyramids for odd lengths and necessity to duplicate degenerate sequences in the even-length group of sequences to achieve antiparallel-complementary duplex pairs.

Application of the C^II^mono constraint drastically reduces the dimensionality of the accessible sequence universe. Whereas the unrestricted Newtonian sequence space is defined by independent variation of four nucleotide coordinates constrained only by total sequence length, the C^II^mono rule collapses this freedom onto a one-dimensional compositional manifold parameterized solely by the relative contribution of A/T and G/C pairs. Geometrically, the C^II^mono filter contracts the unrestricted multinomial sequence universe into a one-dimensional symmetry-constrained subspace. Notably, the number of C^II^mono-compliant sequences grows far more slowly than the total size of the unrestricted sequence universe.

The application of the higher-order C^II^oligo constraint is expected to impose substantially stronger restrictions, as it requires preservation of parity not only for mononucleotide composition but also for all antiparallel-complementary oligonucleotide pairs up to defined k-mer lengths. Consequently, successive application of higher-order symmetry filters progressively partitions the Newtonian sequence universe into increasingly sparse and highly structured subspaces.

Finally, the second Chargaff parity rule implies an increased likelihood of observing **inverted repeats** in genomes. It is therefore noteworthy that **mirror repeats** occur in mouse and human genomes at frequencies comparable to those of inverted repeats, and that the ratio between these two classes of sequence patterns changes during evolution (25,26).

### 2.3 Genome Shadows

In the genomes examined to date, the complete sets of di- and trinucleotides (k-mers of length 2 and 3) are fully represented. As k-mer length increases, however, the fraction of possible k-mers present in any given genome decreases sharply. At the extreme, treating entire chromosomes as single k-mers highlights a fundamental limitation: genome or species identity, as currently defined by a *Reference Genome*, becomes increasingly difficult to reconcile with the full combinatorial sequence space.

Although a full graphical representation of the combinatorial space corresponding to a genome the size of *E. coli* would require far more compact visualization than Figure 1C, the essential point is that this space is structured in the same way as the paired pyramids introduced earlier. The paired pyramids partition the sequence space into symmetry-related halves connected by concerted nucleotide substitutions. The symmetry operation illustrated for tetranucleotides already permits any nucleotide sequence (shown in black in Figure 1C) to generate a derivative sequence (shown in gray) by a concerted substitution in which every A is replaced by G and every T by C. Unlike a conventional mutant, such a derivative differs from the original sequence at every position in a coordinated manner.

In the context of the DNA double helix, each strand is paired with its complementary partner, establishing the familiar C↔G and A↔T relationship defined by the complementarity principle. In addition to this relationship, however, two other pairwise substitution schemes can generate derivative sequences while preserving base pairing within duplexes:

1. Concerted substitutions C↔T and G↔A
2. Concerted substitutions C↔A and G↔T

To illustrate this, consider the sequence 5′-ATGTCA-3′ and its complementary partner 5′-TGACAT-3′. Application of the C↔T/G↔A substitutions yields the sequence pair 5′-GCACTG-3′ and its complement 5′-CAGTGC-3′. Application of the C↔A/G↔T substitutions produces the pair 5′-CGTGAC-3′ and 5′-GTCACG-3′. In all three cases, the original complementarity relationship between strands is preserved.

For clarity, instead of representing these sequences as duplexes, the complementary strands can be appended as continuations of their partners:

1. 5′-ATGTCA–TGACAT-3′
2. 5′-GCACTG–CAGTGC-3′
3. 5′-CGTGAC–GTCACG-3′

The palindromic structure of these illustrative examples is introduced solely to ensure exact compliance with the second Chargaff parity rule within individual strands. If, instead of sequence (1), we consider a full-length chromosomal strand, then sequences (2) and (3) represent two **shadow sequences** of that chromosome. Every genome or chromosome therefore occupies not an isolated position in the Newtonian sequence universe, but a symmetry-related triplet composed of the original sequence and two shadow counterparts. Among the eight 4-mer duplexes that survive application of Rules 1 and 2 (Figure 2C), four constitute examples of such shadow duplexes.

This thought experiment demonstrates that the first two rules—antiparallel-stranded DNA and the second Chargaff parity rule—are **necessary but insufficient** to isolate all biologically compatible sequences from the Newtonian sequence space. Additional shadow transformations can readily be envisaged. For example, a chromosome may acquire a shadow by inversion of its 5′–3′ polarity, or by systematic permutation of adjacent bases, such as transforming 1234 into 2143 across the entire sequence.

The symmetry operations described above imply that biologically compatible sequences do not occupy isolated positions within the Newtonian sequence universe. Instead, they form interconnected families related by complementarity, reversal, shadow substitutions, and higher-order transformations. For longer sequences, the natural objects of the filtered sequence space therefore become symmetry-related sequence classes, or orbits, rather than individual nucleotide strings. For a sequence *S*, an orbit may be represented schematically as the set of sequences generated through repeated application of symmetry operations such as reversal (*R*), complementarity (*C*), reverse complementarity (*RC*), and shadow transformations.

## Summary of the Three Filtering Rules

The Newtonian sequence space defined by the multinomial expansion of four nucleotides contains an overwhelming number of sequences that are biologically irrelevant. The reduction of this space toward biologically compatible sequences can be formalized as a hierarchy of three filters, each eliminating entire symmetry classes of sequences by invoking fundamental constraints imposed by DNA structure and genome organization.

## Rule 1: Antiparallel-stranded duplex DNA with A–T and G–C base pairing

The requirement that genetic material be replicated and maintained as an antiparallel double helix immediately excludes a large class of sequences that are compatible with parallel-stranded pairing. Parallel complementarity permits strand self-crossover and Möbius-type topologies, favoring recombination and sequence reshuffling rather than faithful preservation. Sequences composed of two halves related by parallel complementarity are therefore disfavored as templates for stable hereditary information storage. This rule imposes the first, structurally grounded reduction of sequence space.

## Rule 2: The Second Chargaff Parity Rule (SCPR)

The second Chargaff parity rule further constrains sequence space by requiring approximate equality of A and T, and of G and C, within each single strand (C^II^mono), and, at higher resolution, equality of antiparallel-complementary k-mers (C^II^oligo). Exact C^II^mono symmetry is restricted to even-length sequences, whereas odd-length sequences satisfy only parity-limited approximations of the rule. Application of this rule eliminates sequences lacking sufficient compositional balance, particularly at short lengths, and drastically reduces the number of admissible sequences. SCPR reflects long-term equilibrium constraints imposed by replication, mutation, and repair, and acts as a powerful statistical filter on sequence space.

## Rule 3: Genome Shadows

Even after application of Rules 1 and 2, additional sequences remain that satisfy structural and compositional constraints but are related to real genomes by global symmetry transformations. These *genome shadows* define symmetry-related counterparts of real genomes generated through concerted nucleotide substitutions, strand polarity inversion, or systematic positional permutations, all of which preserve duplex complementarity and, in some cases, SCPR. The existence of such shadows demonstrates that Rules 1 and 2 are necessary but insufficient to uniquely specify biologically relevant genomes. Rule 3 introduces a higher-order constraint, acknowledging that only a limited subset of symmetry-related sequences can function as whole genomes, while others are progressively eliminated as biologically incompatible.

Together, these three rules define a framework in which biologically meaningful sequences emerge as a small, symmetry-restricted subset of the Newtonian sequence space. In this framework, genome architecture is shaped not only by local sequence motifs but by global symmetry breaking, linking sequence composition, topology, and evolutionary dynamics.

### 3.1 Sequence Orbits in the Filtered Space

Application of the three filtering rules reduces the Newtonian sequence universe to a highly constrained subset. The surviving sequences are not distributed uniformly throughout this reduced space. Instead, they form groups of related sequences connected by symmetry operations such as duplex complementarity, shadow transformations, strand reversal, and combinations thereof. In the present work, such groups are referred to as sequence orbits. The term orbit is used in the group-theoretical sense to denote the set of sequences generated from a given sequence by application of the allowed symmetry transformations and does not imply an evolutionary trajectory.

The tetranucleotide duplexes shown in Figure 1C already provide a simple example. Each duplex is accompanied by its shadow counterpart generated through concerted A↔G and T↔C substitutions. Together with duplex complementarity, these transformations partition the sequence space into families of related sequences. The same organization persists at larger sequence lengths, although the number of members and the complexity of the resulting orbit structures increase rapidly.

The emergence of orbit structure follows naturally from the filtering process itself. Rule 1 removes sequences compatible with parallel-complementary strand organization. Rule 2 restricts the surviving sequences to a parity-constrained compositional manifold defined by the second Chargaff parity rule. Rule 3 identifies additional symmetry-related shadow counterparts generated through global sequence transformations. As a consequence, biologically meaningful sequences no longer appear as isolated points in sequence space but as members of larger symmetry-related families.

This distinction is particularly important because the filters do not simply reduce the number of admissible sequences. They also impose internal organization upon the surviving subset. Sequences that satisfy the same compositional constraints may nevertheless occupy different orbit families and therefore possess different symmetry properties.

The orbit concept provides a natural language for describing the filtered sequence universe. Rather than viewing genomes as isolated strings of nucleotides, genomes may be regarded as representatives of symmetry-related classes embedded within the larger Newtonian sequence space. As sequence length increases, the number of possible sequences grows exponentially, whereas the fraction surviving the filters becomes progressively smaller. The biologically relevant problem therefore shifts from enumeration of sequences to characterization of the orbit families that remain.

Within this framework, evolutionary processes such as mutation, recombination, insertion, deletion, duplication, and horizontal transfer may be interpreted as movements between neighboring regions of the filtered sequence universe. The filters define the admissible space; the orbit structure defines its internal organization.

The persistence of simple sequence motifs after application of the filters is particularly noteworthy. Examination of SCPR-compatible sequences reveals that the surviving subset is not composed exclusively of fully unique arrangements of nucleotides. Mirror-symmetric sequences, reverse-complement palindromes, repetitive motifs, and other internally ordered patterns continue to occur. These patterns do not represent independent filtering rules. Rather, they correspond to special positions within the orbit structure and therefore provide insight into the residual symmetry retained by the filtered sequence space.

### 3.2 Orbit Degeneracy and Symmetry

Most sequences generate large orbit families in which each symmetry operation produces a distinct sequence. Certain sequences, however, possess internal symmetries that cause multiple transformations to yield identical outcomes. Such sequences occupy reduced orbits and therefore represent singular points within the filtered sequence space.

Several classes of symmetry-fixed sequences can be distinguished. Mirror-symmetric sequences satisfy

S = R(S)

where R denotes reversal of sequence order. In general form, such sequences may be represented as

abcd|dcba

for even lengths, or abcdXdcba

for odd lengths. Because reversal leaves these sequences unchanged, they occupy reduced orbit positions.

A distinct class consists of reverse-complement palindromes satisfying S = RC(S)

where RC denotes the reverse-complement operation. These are the classical palindromic sequences encountered in duplex DNA. Such sequences are invariant under reverse complementarity rather than simple reversal. Consequently, mirror symmetry and reverse-complement symmetry define related but distinct classes of orbit degeneracy.

Periodic and repetitive sequences constitute another class of reduced-orbit objects. These sequences contain repeated sequence units and therefore remain invariant under selected translational or periodic transformations. Although simple periodic sequences are strongly restricted by the filtering rules, periodicity persists within many SCPR-compatible sequence families.

Finally, some sequences exhibit self-shadowing behavior. In such cases, application of one or more shadow transformations produces either the original sequence or another sequence belonging to the same reduced orbit. Exact self-shadowing fixed points are expected to be rare, but partial self-shadowing becomes increasingly important as sequence length increases and multiple symmetry operations act simultaneously.

These classes do not define additional filtering rules. Rather, they represent symmetry-fixed subsets embedded within larger orbit families. Their persistence after application of the first three filters demonstrates that biologically admissible sequence space retains substantial internal organization and is not composed exclusively of fully unique sequences.

The existence of reduced orbits further suggests that the filtered sequence universe possesses a hierarchical internal structure. Large generic orbits coexist with smaller symmetry-constrained orbit families, analogous to general and special positions in crystallographic space groups. Characterizing these orbit classes may therefore provide a useful framework for describing the organization of biologically compatible sequence space.

The sequence-space framework developed above identifies a variety of symmetry relationships that persist after application of the three filtering rules. These include mirror symmetry, reverse-complement symmetry, shadow transformations, and relationships between parallel- and antiparallel-complementary sequence arrangements. The persistence of such motifs demonstrates that biologically admissible sequence space retains substantial internal organization even after strong structural and compositional constraints have been imposed.

Among these surviving patterns, mirror-symmetric sequences are of particular interest because they occupy an intermediate position between conventional antiparallel duplex DNA and alternative pairing arrangements. Their existence raises a natural question: can the symmetry relationships identified within the filtered sequence space be realized physically by known nucleic-acid structures?

Importantly, the relationship between parallel-stranded and antiparallel-stranded sequences is not one of mutual exclusivity(17).

Figure 2C illustrates a circular double-stranded DNA molecule in which one half of the sequence is mirror-symmetric. This segment reads as a palindrome, here emphasized by rendering a classical Greek text on the top strand to distinguish mirror symmetry from conventional palindromes defined on antiparallel duplexes.

This circular construct can be topologically transformed into a junction between parallel-stranded and antiparallel-stranded DNA segments connected at a crossing point (Figure 2C, bottom panel, left). If the two strands of the parallel-stranded circle are cut at the position marked by the black arrow and reconnected after a 180° rotation—analogous to the transformation shown in Figure 2B—the resulting structure assumes a single-stranded Möbius-strip topology (Figure 2C, bottom panel, right). This topological feature suggests that mirror-symmetric segments may serve as crossing points at which intrastrand recombination can occur.

The structural implications of such mirror-symmetric arrangements extend beyond the duplex itself. The equilibrium between parallel- and antiparallel-aligned segments with mirror-symmetric complementarity depends on the relative stability of the secondary structures adopted by these conformations. A junction between trans-paired and cis-paired duplexes cannot be accommodated without local strand separation unless the structure is realized as a G-quadruplex, in which cis- and trans-base-pair-aligned elements coexist within stacked tetrads (Figure 2D, right panel). Indeed, the intersection highlighted by the dashed rectangle in Figure 2C (bottom left) possesses the connectivity expected for a hybrid quadruplex core and illustrates how four duplex segments can be reorganized into a quadruplex-like junction.

This demonstrates how mirror symmetry can be propagated into higher-order DNA junctions. Among currently known DNA conformations, G-quadruplexes occupy a unique position because they combine structural features that are otherwise distributed among several nucleic-acid architectures, including parallel and antiparallel strand orientations, cis- and trans-like pairing relationships, strand-crossing topologies, and junctions between multiple DNA segments. G-tetrads form stable secondary structures composed of four guanines arranged in a planar array, stabilized by hydrogen bonding involving Watson–Crick and Hoogsteen edges and coordinated by monovalent cations that neutralize the negative charge of the O6 oxygens (Figure 2D, right panel). Stacked G-tetrads form G-quadruplex DNA. From one perspective, a G-quadruplex can be viewed as an associate of two duplexes containing cis base pairs that are isomorphic to Watson–Crick base pairs. In this framework, a guanine paired through its Hoogsteen edge (α-guanine) is functionally equivalent to a pyrimidine, whereas a guanine paired through its Watson–Crick edge (β-guanine) is equivalent to a purine (Figure 2D), consistent with the base-pairing classification of Leontis and Westhof(19).

Each guanine in a G-tetrad may adopt syn or anti glycosidic conformations, allowing adjacent guanines with anti/syn combinations to transition smoothly into anti/anti Watson–Crick base pairs. This continuity is illustrated in Figure 2D by the alignment between α/β guanines and a C·G base pair. Moreover, two diagonal guanines within a tetrad can be extended into a trans base pair. Trans A·T base pairs are known to play this structural role, including 5′–5′ interactions (27) and 3′–5′ interactions with diagonal guanines in a G-tetrad (28). Purine–purine trans pairs are particularly well suited to span quadruplex diagonals, and trans A·A Watson–Crick base pairs have been observed at the 3′ ends of dimeric hexads (29), in the c-MYC quadruplex (30), and at the interface connecting two quadruplex halves in the dimeric c-KIT2 structure (28). Two successive isomorphic trans A·A and G·G base pairs may even form a parallel-stranded mini-helix within a quadruplex(31).

A third perspective views the G-tetrad as an associate of a guanine triplet plus a single base. Since triplex nucleic acids frequently involve a third strand forming hydrogen bonds with the Hoogsteen edge of a Watson–Crick duplex, the position of this third base is isomorphic to that of the fourth guanine in a tetrad. Consistent with this view, guanine triplets with a vacant fourth position are known constituents of G-quadruplexes(32,33), and aptamers containing quadruplex–triplex junctions have been described (34). A genome-wide bioinformatic analysis of E. coli identified recurring motifs capable of alternatively adopting triplex or quadruplex conformations (35). Soon after their discovery at telomeric ends, G-quadruplexes were proposed as intermediates facilitating DNA rearrangements during meiosis and immunoglobulin class switching (12,13). Subsequent genome-wide analyses revealed that G-rich sequences are strongly associated with elevated recombination rates and genome rearrangements. Triple-helical DNA structures have likewise been implicated as intermediates in homologous recombination.

Taken together, these observations demonstrate that structural elements exist which naturally combine symmetry relationships that appear separately in duplex and triplex DNA. In this regard, the G-quadruplex occupies a distinctive position among known nucleic-acid structures. Rather than representing merely another secondary-structure motif, it provides a structural framework capable of accommodating relationships otherwise distributed among distinct nucleic-acid conformations, including cis and trans base pairing, parallel and antiparallel strand orientations, and strand-crossing topologies. These are precisely the classes of symmetry relationships identified in the filtered sequence space. The G-quadruplex may therefore be viewed as a potential physical intermediary between symmetry classes that otherwise appear disconnected in the combinatorial description of sequence space.

If G-quadruplexes indeed provide structural intermediates capable of linking otherwise distinct symmetry relationships, one would expect at least some biological systems to exploit these properties during DNA rearrangement. One of the best characterized examples is the RecA-dependent pilE antigenic variation system of *Neisseria gonorrhoeae*, in which a parallel-stranded G-quadruplex has been shown to be essential for homologous recombination.

### 4.2 RecA and the pilE Quadruplex

Direct evidence for the functional involvement of mirror-symmetric DNA segments in genome rearrangement has been obtained in studies of pilin antigenic variation in Neisseria gonorrhoeae (36). There are strong reasons to suspect that related principles operate in other, less extensively characterized bacterial systems(37).

The *pilE* G-quadruplex has long been recognized as an essential element of pilin antigenic variation in *N. gonorrhoeae*. Following determination of the solution structures of the monomeric and dimeric *pilE* G-quadruplexes, we proposed that the compact three-layered monomeric quadruplex could be accommodated between the two L2 loops of adjacent RecA monomers within the active filament (38). At that time, however, neither high-resolution structures of RecA-mediated strand exchange nor the biological importance of quadruplex loop architecture had been established.

Subsequent genetic and biochemical studies demonstrated that the biological activity of the *pilE* quadruplex depends critically on RecA and on the size and composition of the quadruplex loops(36,39), while recent structural studies have provided a much more detailed picture of the RecA filament during homologous recombination. Independent biochemical evidence likewise indicated that RecA distinguishes among parallel-stranded G-quadruplexes according to their structural architecture. Paul et al. (40) showed that RecA readily unwinds a telomeric parallel-stranded G-quadruplex containing three-nucleotide propeller loops, whereas the three-layered promoter quadruplex from c-MYC remained compact and resistant to unwinding under otherwise similar conditions. Together with the pilE genetic studies, these observations suggested that productive interaction with RecA depends not merely on quadruplex formation, but on specific geometric features of the quadruplex architecture. These advances now permit a comprehensive structural re-evaluation of the original hypothesis.

Accordingly, we examined a series of G-quadruplex architectures differing in tetrad number, loop organization, and sequence composition in order to determine the structural features required for productive interaction with the RecA filament and to place the native *pilE* quadruplex within this broader structural landscape.

#### 4.2.1 Structural Compatibility of G-Quadruplexes with RecA

##### Four-layered, four-stranded all-parallel G-quadruplex

In vivo replacement of the native pilE G-quadruplex with four-layered quadruplex variants abolishes pilin antigenic variation(36). Consistent with this observation, our structural modeling and computational analysis shows that four-layered all-parallel G-quadruplexes are generally incompatible with the geometry of the active RecA filament.

Across the ensemble of top-scoring docking solutions, the four-layered quadruplex core is positioned away from the inter-Loop 2 region of the filament, which is implicated in strand exchange. Instead, these quadruplexes preferentially bind within a surface cavity formed by the β6/β7 hairpin, the associated β8 strand, and the C-terminal domain (CTD) of RecA (Figure 3A,B; (see (41)for nomenclature)). Two binding orientations are observed, involving either the 5′- or 3′-terminal tetrad facing the protein surface (figure 3 A and B).

**Figure 3.**
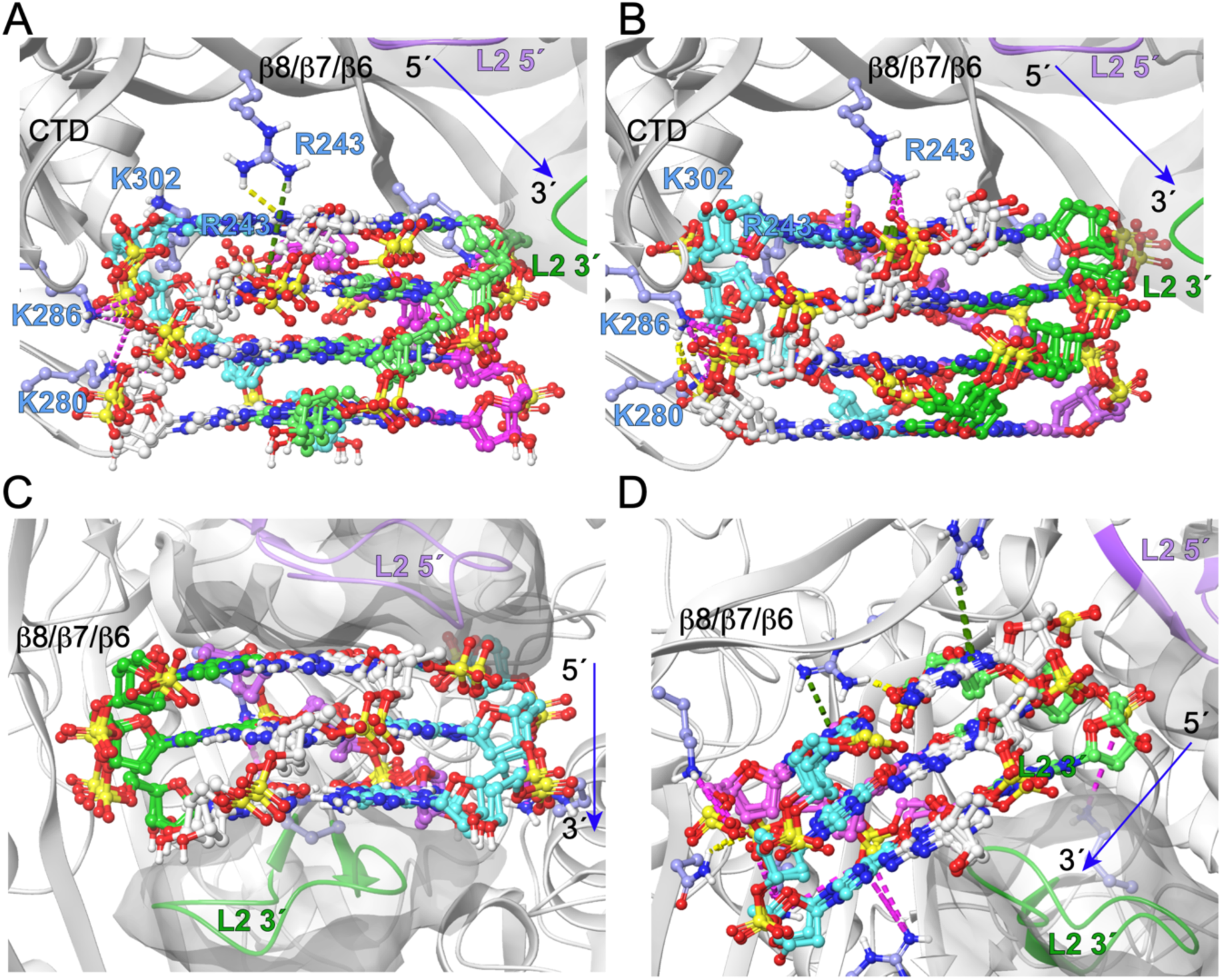
Structural compatibility of loop-free all-parallel G-quadruplexes with the RecA filament. Results of docking calculations for four-layered (A,B) and three-layered (C,D) all-parallel G-quadruplexes composed of four independent guanine strands lacking connecting loops. The two L2 loops that define the inter-L2 region of adjacent RecA monomers are shown as ribbon/surface representations, with the 5′-side Loop 2 colored lilac and the 3′-side Loop 2 colored green. The blue arrow indicates the 5′→3′ direction and inclination of the RecA filament axis. **(A,B)** Four-layered G-quadruplexes reproducibly bind within the cavity formed by the C-terminal domain (CTD) and the inter-L2 region rather than between the two L2 loops. Two principal binding orientations are observed, with either the 5′-terminal tetrad (A) or the 3′-terminal tetrad (B) facing the protein surface. Stabilization is mediated primarily by interactions involving Lys280, Lys286, Lys302, Arg243, and the phosphate backbone of the quadruplex. **(C,D)** Three-layered G-quadruplexes exhibit productive interaction modes compatible with the active RecA filament. In the highest-scoring poses, the quadruplex is either fully sandwiched between the two L2 loops (C) or partially inserted into the inter-L2 region while maintaining favorable interactions with both adjacent RecA monomers (D).

Stabilization is mediated predominantly by electrostatic interactions between basic RecA residues and the quadruplex phosphate backbone. A prominent and reproducible feature in all binding modes is interaction of K280, K286 and K302 with the phosphate backbones and groove formed by strands 1 and 4. R243 further contributes through π–cation interactions with guanine bases and salt bridges to adjacent phosphate groups, while additional lysine residues engage the phosphate backbone in the remaining grooves Figure 3 A and B).

Importantly, in none of the observed configurations does the four-layered quadruplex fit between the two Loop 2 regions of adjacent RecA monomers, providing a structural explanation for its inability to support antigenic variation in vivo.

##### Three-layered, four-stranded all-parallel G-quadruplex

In contrast to the four-layered topology, three-layered all-parallel G-quadruplexes exhibit interaction modes with RecA that are fully compatible with the geometry of the active filament. Among the top-ranked docking solutions, several configurations position the quadruplex core directly between the Loop 2 regions of adjacent RecA monomers, where favorable van der Waals complementarity stabilizes the complex Figure 3 C.

In these inter-Loop 2 binding modes, phosphate groups from two opposing strands of the quadruplex form salt bridges with basic residues from neighboring RecA monomers, effectively bridging the filament across its functional interface. This results in directed, symmetric interactions involving only a subset of quadruplex strands, while the remaining strands face away from the protein surface.

Additional high-scoring poses place the three-layered quadruplex within Site II of the RecA filament, with the quadruplex axis inclined across two Loop 2 regions. In this configuration, one strand engages multiple positively charged residues from two adjacent RecA monomers through electrostatic interactions and π–cation contacts involving terminal guanine bases. A neighboring strand is partially inserted between the Loop 2 regions, further stabilizing the complex via phosphate-mediated interactions, whereas the remaining strands contribute minimally or not at all to protein binding Figure 3 D.

Importantly, all productive interaction modes observed for the three-layered quadruplex preserve accessibility of the inter-Loop 2 space and maintain compatibility with RecA-mediated strand exchange, providing a structural rationale for the ability of three-layered G-quadruplexes to support pilin antigenic variation in vivo.

##### Three-layered parallel-stranded G-quadruplexes with extended propeller loops

Replacement of the native pilE G-quadruplex with other G-rich sequences from the N. gonorrhoeae genome (IGR1172 and NGO816), both capable of forming three-layered G-quadruplexes, results in the loss of pilin antigenic variation (36). In their all-parallel conformations, these sequences form G-quadruplexes with three-nucleotide propeller loops extruding outward from the core, resembling the crystallographically characterized human telomeric parallel-stranded G-quadruplex (42). This topology was therefore used to probe the effect of enlarged loops on RecA–quadruplex compatibility.

Docking calculations reveal several distinct binding modes for this loop-containing three-layered quadruplex; however, none reproduce the inter–Loop 2 sandwiching observed for the loop-free pilE quadruplex. In a minority of poses, the quadruplex core is positioned between Loop 2 regions of adjacent RecA monomers, but the orientation of its DNA strands is inverted relative to both the pilE quadruplex and the RecA-bound ssDNA (Figure S1 A). Moreover, accommodation of the bulky propeller loops displaces the core outward from the filament axis, reducing geometric complementarity with the recombination-active interface (Figure S1 B).

In other prevalent binding modes, the quadruplex associates with peripheral regions of the RecA filament, including the β6/β7 hairpin and the C-terminal domain, similarly to the behavior observed for four-layered quadruplexes (Figure S1 C and D). In these configurations, the presence of large loops forces a tilt and lateral shift of the quadruplex core, further precluding stable insertion between Loop 2 regions.

Additional poses involve interactions mediated primarily by contacts with the propeller loops themselves (Figure S1 E) or by auxiliary helices from neighboring RecA monomers (Figure S1 F). These interactions stabilize the complex but do not position the quadruplex in a manner compatible with RecA-mediated strand exchange.

Taken together, these results indicate that while a three-layered parallel-stranded G-quadruplex core is compatible with RecA in principle, the presence of large propeller loops disrupts the precise geometric and orientational requirements necessary for productive engagement with the recombination-active region of the filament. The loops do not appear to prevent binding directly; rather, they alter the orientation and accessibility of the quadruplex core required for productive engagement with the inter-Loop 2 region. This provides a structural explanation for the inability of telomeric-like and pilE-replacement quadruplexes with enlarged loops to support antigenic variation in Neisseria Gonorrhoeae in vivo.

#### 4.2.2 Structural Basis of pilE Quadruplex Recognition

##### PilE monomeric G-quadruplex

The native pilE G-quadruplex possesses an unusually compact loop architecture that distinguishes it from other known parallel-stranded three-layered quadruplexes. In the all-parallel monomeric pilE G-quadruplex, three propeller loops connect adjacent guanine tetrads across three of the four grooves. Two of these loops are single-residue thymidines (dT4 and dT13), while the third loop (dT8–dT9) comprises two residues but adopts an unusually compact conformation due to inward orientation of dT9. This inward-directed base is stabilized by hydrogen bonding between O2 and H3 of dT9 and the N3 and H22 atoms of guanines dG6 and dG7 in the middle and 3′-terminal tetrads, respectively, thereby effectively reducing the exposed size of the propeller loop (38). Viewed along the quadruplex axis (Figure S2), the three compact propeller loops occupy distinct sectors around the quadruplex circumference, creating an asymmetric recognition surface that can engage complementary regions of adjacent RecA monomers. Docking calculations show that all productive interaction modes of the pilE quadruplex position its core between the Loop 2 regions of two adjacent RecA monomers, consistent with the geometry of the active RecA filament (43). Unlike loop-free three-layered quadruplexes, whose docking orientations remain relatively degenerate, the compact propeller loops of the pilE quadruplex uniquely define the orientation and register of the G-quadruplex within the RecA filament (Figure 4 A and B).

**Figure 4.**
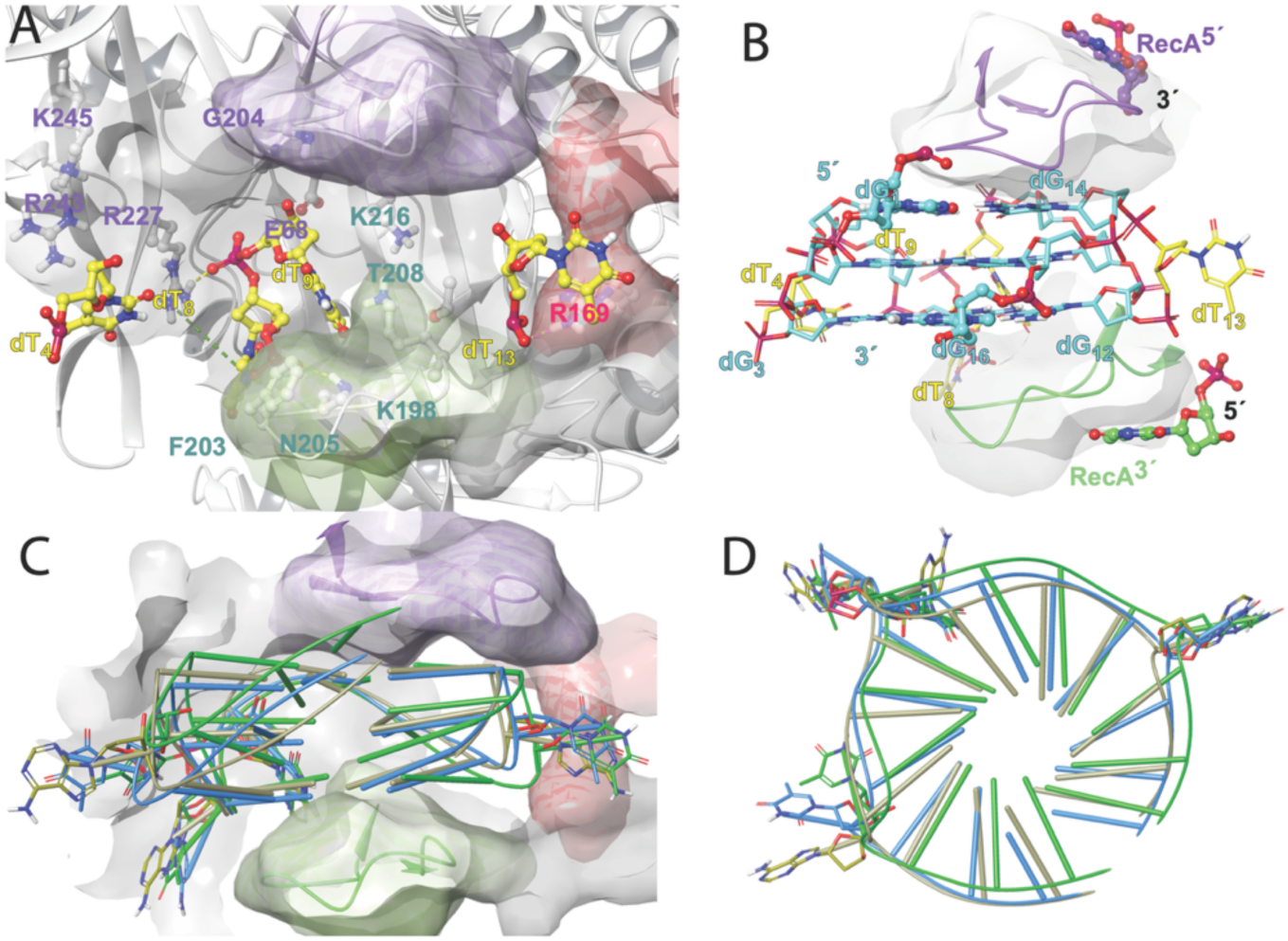
Structural basis for recognition of the pilE G-quadruplex by the RecA filament. **(A)** Docking calculations show that the native pilE G-quadruplex reproducibly inserts between the L2 loops of adjacent RecA monomers. The 5′-side and 3′-side Loop 2 regions are colored lilac and green, respectively. Guanine residues of the quadruplex core are omitted to emphasize the positions of the dT4, dT8–dT9, and dT13 propeller loops, which reproducibly interact with the indicated RecA residues. **(B)** The complete pilE G-quadruplex shown in the context of neighboring RecA inter-L2 binding sites. The adjacent 5′- and 3′-terminal trinucleotide positions of the RecA filament are indicated schematically. Note that neither the 5′-terminal residue (dG1) nor the 3′-terminal residue (dG16) can be directly connected to the neighboring RecA-bound ssDNA without local structural adjustment. **(C,D)** Docking models of antigenic variation (AV)-active pilE G-quadruplex variants containing T–TT–AT (blue) and A–AA–A (olive) loop combinations, superimposed with the native pilE quadruplex (cyan). Panel **C** shows the docked complexes within the RecA filament, whereas panel **D** presents top views of the superimposed quadruplex ribbon representations. Despite differences in loop composition, the compact loop architectures adopt highly similar spatial positions, preserving productive interactions with the RecA filament.

In all observed poses, the dT4 and dT8–dT9 loops interact directly with a continuous positively charged surface formed by residues Arg227, Arg243, and Lys245 of the ATP-binding core of one RecA monomer, as well as Lys216 from helix αG and Thr208 from Loop 2 of the adjacent monomer (Figure 4A). The dT13 loop engages a spatially distinct surface defined by residues from Loop 1, including the M164–R169 segment, which has been implicated in discriminating between single-stranded and double-stranded DNA binding by the RecA filament (Figure 4A)(44,45). Remarkably, the sixteen-nucleotide pilE G-quadruplex fits entirely within the space between two Loop 2 regions of the RecA filament, a region that normally accommodates a three-nucleotide recombination unit. The orientation of the quadruplex strands coincides with that of the filament-forming single-stranded DNA, providing a structural explanation for why the native pilE quadruplex, unlike other examined parallel-stranded G-quadruplexes, is uniquely compatible with RecA-mediated homologous recombination.

##### N. gonorrhoeae pilE quadruplex derivatives with variable loop size and composition

Recent in vivo and in vitro studies have demonstrated that modest alterations in the size and composition of pilE G-quadruplex loops have pronounced effects on pilin antigenic variation and RecA binding affinity (39). Replacement of the native T:TT:T loop combination with either T:TT:AT or A:AA:A preserved antigenic variation, albeit at reduced levels, whereas substitution with AT:ATT:AT or TTT:GC:ATC resulted in complete loss of the phenotype and diminished RecA affinity. The docking results provide a structural rationale for these observations. As in the native *pilE* quadruplex, the second thymidine of the AT loop can adopt an inward-directed conformation, effectively reducing the functional loop size to a pseudo-single-nucleotide loop. Consequently, quadruplexes containing T:TT:AT or A:AA:A loop combinations predominantly adopt poses that fit within the inter-Loop 2 region of the RecA filament (Figure 4 C and D), closely resembling the native *pilE* quadruplex. As in the native pilE quadruplex, the second thymidine in the AT loop can adopt an inward-directed conformation, effectively rendering the loop pseudo-single-residue in size. Accordingly, quadruplexes containing T:TT:AT or A:AA:A loop combinations predominantly adopt poses that fit within the inter–Loop 2 space of the RecA filament, similar to the native pilE structure. In contrast, loop combinations incapable of adopting such compact conformations behave similarly to telomeric-like three-layered quadruplexes containing extended propeller loops, leading to displacement or incorrect orientation of the quadruplex core relative to the recombination-active region of the filament. Thus, the computational models closely parallel the experimental phenotype: variants that preserve compact loop geometry retain productive RecA recognition and antigenic variation, whereas variants with enlarged loops fail to achieve the docking geometry required for engagement of the inter-Loop 2 interface.

#### 4.2.3 Integration of the pilE Quadruplex into the RecA Filament

##### 3’- and 5’-extension of the PilE quadruplex into RecA filament

The RecA filament possesses an approximately sixfold helical symmetry, with successive RecA monomers related by ∼60° rotation around the filament axis. The bound ssDNA follows the same periodicity, with successive trinucleotide units adopting corresponding rotational increments (Figure 2A). In contrast, superposition of the 5′- and 3′-terminal tetrads of the *pilE* G-quadruplex reveals that the quadruplex itself possesses an intrinsic negative rotational offset of approximately 15° between its 5′ and 3′ ends (Figure 5B).

**Figure 5.**
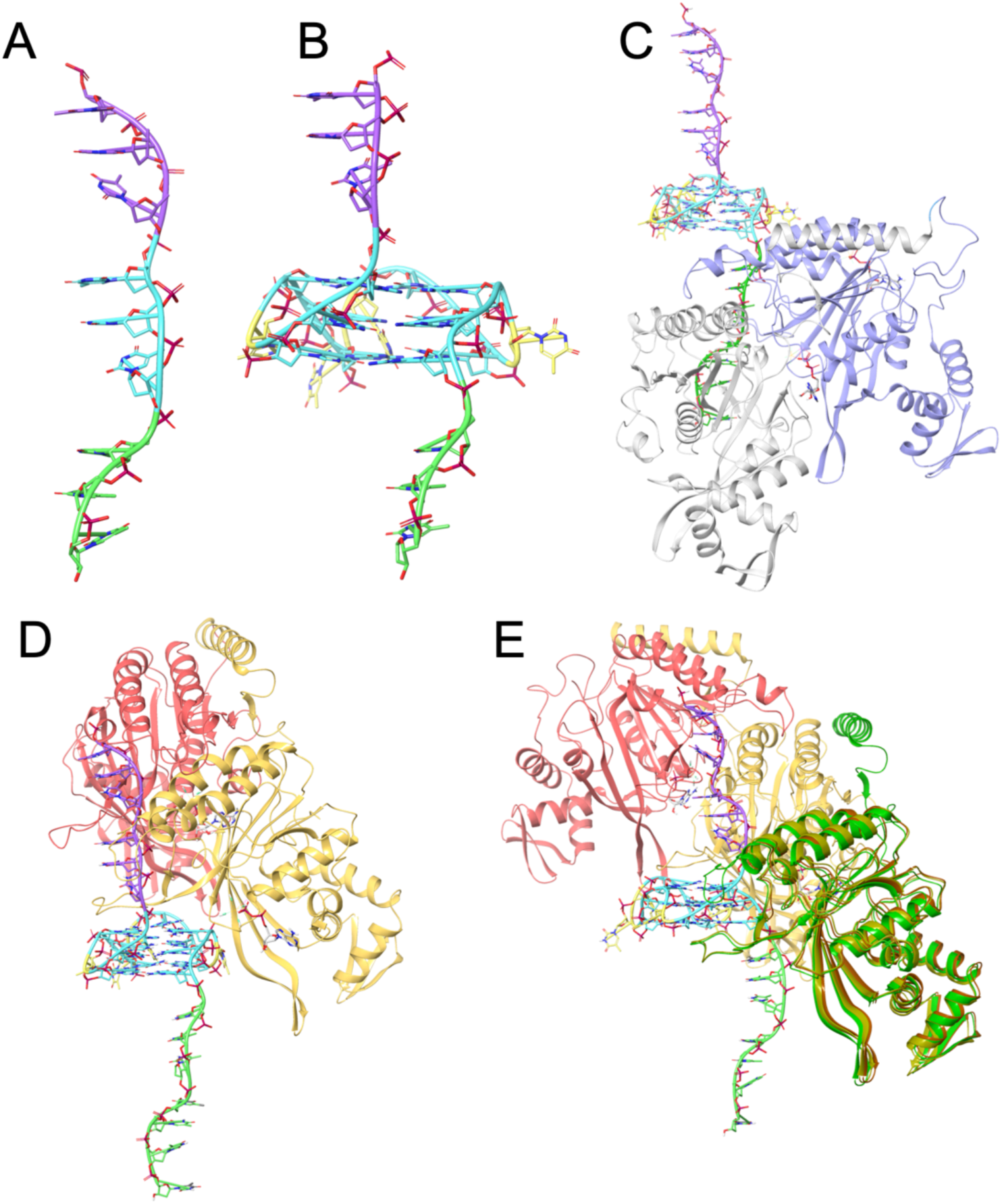
Integration of the pilE G-quadruplex into the RecA filament and directional filament assembly. **(A,B)** Comparison of RecA-bound ssDNA and the pilE G-quadruplex. **(A)** A 9-nucleotide ssDNA in the RecA filament conformation, with the central trinucleotide colored cyan, the 5′-terminal trinucleotide lilac, and the 3′-terminal trinucleotide green. **(B)** The pilE G-quadruplex (cyan) shown with corresponding 5′-(lilac) and 3′-terminal (green) trinucleotide extensions. Note that, unlike RecA-bound ssDNA, the quadruplex imposes a negative rotational relationship between the 5′- and 3′-terminal extensions. **(C,D)** Models of the pilE G-quadruplex extended by six nucleotides of ssDNA in the RecA filament conformation. **(C)** Two RecA monomers bound to the 3′-terminal ssDNA extension. **(D)** Two RecA monomers bound to the 5′-terminal ssDNA extension. The two extension modes differ by rotation of the quadruplex core by one guanine column (see Figure S2). **(E)** Directionality of RecA filament assembly. Only the complex containing the 5′-terminal ssDNA extension (**D**) supports productive docking of an additional RecA monomer adjacent to the quadruplex. In the resulting complex, the inserted L2 loop is positioned beneath the 3′-terminal tetrad, and the docked monomer (green) closely superimposes on the corresponding crystallographic RecA monomer (olive), demonstrating compatibility with continued filament growth.

This geometric mismatch raises an important structural question: can a folded G-quadruplex be incorporated into the otherwise regular RecA filament while maintaining continuity of the filament-bound ssDNA?

To address this question, models were constructed in which the *pilE* quadruplex was extended by RecA-compatible trinucleotide segments either at its 5′ end or at its 3′ end (Figure S2 A and B). In both cases, the added trinucleotide aligned continuously with the crystallographic ssDNA trajectory, allowing the quadruplex-containing DNA to occupy successive inter-Loop 2 sites within the RecA filament (Figure S2 C and D).

Although both extensions are geometrically compatible with the filament, they differ in the orientation of the quadruplex core. Relative to the 5′-extended form, the 3′-extended quadruplex undergoes an approximately one-tetrad rotational shift (Figure S2 E and F).

This rotation changes the protein-facing surface of the quadruplex while preserving continuity of the attached ssDNA.

In the 3′-extended model, loops dT4 and dT8–dT9 remain solvent exposed, while the compact dT13 loop becomes oriented toward the protein, where its single-residue geometry allows close packing without significant steric clashes (Figure 6 A and B).

**Figure 6.**
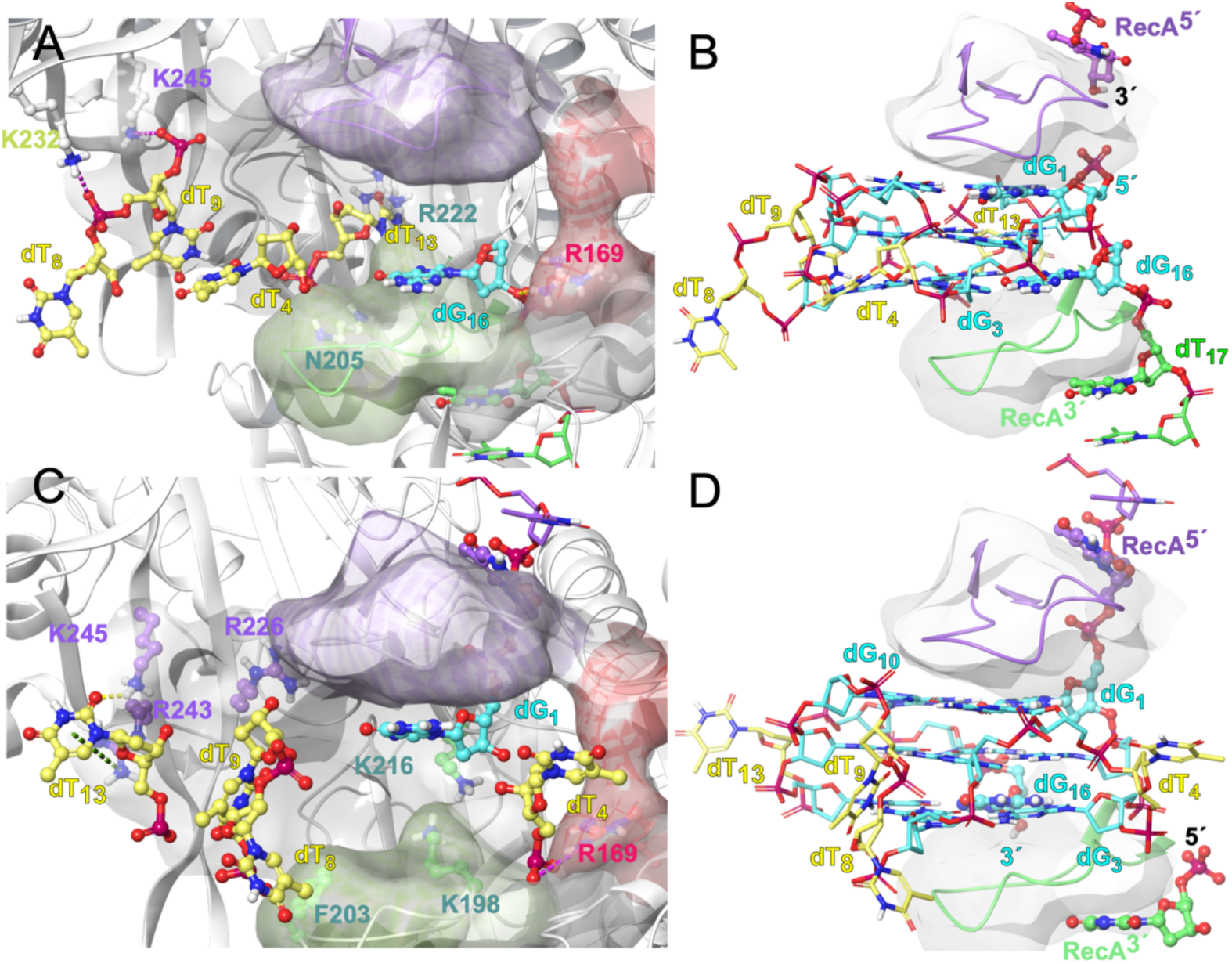
One-column rotation of the pilE G-quadruplex associated with alternative ssDNA extension modes. **(A,B)** Docking model of the pilE G-quadruplex carrying a 3′-terminal ssDNA extension. **(A)** The complex is shown in the context of the complete RecA monomers, with guanine residues omitted to emphasize the positions of the compact propeller loops relative to the protein surface. **(B)** The same docked complex shown as the complete G-quadruplex positioned between the L2 loops of adjacent RecA monomers, represented as ribbons and semi-transparent molecular surfaces. **(C,D)** Corresponding docking model of the pilE G-quadruplex carrying a 5′-terminal ssDNA extension. **(C)** The RecA–quadruplex complex displayed as in panel A to highlight the orientation of the propeller loops following one-column rotation of the quadruplex core. **(D)** The complete docked quadruplex shown between the L2 loops as in panel B. Comparison of the two extension modes demonstrates that extension from opposite termini is accompanied by rotation of the quadruplex core by one guanine column (see Figure S2), resulting in different presentations of the grooves and propeller loops to the RecA filament while preserving accommodation of the compact pilE G-quadruplex within the inter-L2 binding site.

In the 5′-extended model, the 5′–3′ groove faces the protein surface, whereas loops dT4, dT8–dT9 and dT13 project toward solvent. (Figure 6 C and D).

In both cases, the six-nucleotide extensions beyond the quadruplex coincide closely with the conformation of filament-bound ssDNA observed crystallographically.

The one-tetrad rotation relating the two extension geometries is equivalent to the rotational transformation previously described for the triplex-mediated strand-crossing transition between parallel and antiparallel DNA arrangements (Section 4.1), providing an additional structural link between triplex and quadruplex recombination intermediates.

##### Assembly of the RecA filament on quadruplex-containing DNA

The preceding calculations assume that the quadruplex is already embedded within an assembled RecA filament. An equally important question is whether filament assembly itself can propagate across a folded *pilE* quadruplex.

To examine this possibility, partial RecA assemblies were constructed in which a protein-free *pilE* quadruplex was flanked by short ssDNA segments preorganized in the RecA conformation (Figure 5B). RecA monomers were then docked either onto the 5′ side of the quadruplex (Figure 5C) or onto its 5′ side (Figure 5D), allowing filament growth to be examined in both directions.

The docking calculations revealed a pronounced directional asymmetry. Assembly progressing in the physiological 5′→3′ direction generated RecA monomer positions that closely reproduced those observed in crystallographic RecA filaments, including correct placement of the Loop 2 region beneath the terminal tetrad (Figure 5E). By contrast, propagation in the opposite direction failed to generate crystallographically consistent assemblies.

These calculations therefore support a structurally self-consistent model in which a folded *pilE* quadruplex can be incorporated into a growing RecA filament while preserving the overall architecture of the filament.

##### Summary of G-quadruplex–RecA compatibility

Taken together, the docking calculations define a remarkably narrow structural window for productive RecA recognition. Four-layered quadruplexes fail because their cores cannot be accommodated within the recombination-active region of the filament. Three-layered quadruplexes possessing extended propeller loops retain the correct core geometry but lose the precise orientation required for productive inter-Loop 2 binding. The native *pilE* quadruplex combines a three-layered core with compact inward-directed loops that uniquely satisfy both requirements, thereby providing a structural explanation for its exceptional biological activity.

More broadly, they demonstrate that only a highly restricted subset of mirror-symmetric quadruplex structures remains compatible with RecA-mediated genome rearrangement, providing a direct biological example of how structural constraints carve functional trajectories through the filtered sequence space introduced above.

### 4.3 A Structural Model for pilE Gene Conversion Biological context

Antigenic variation in *N. gonorrhoeae* occurs through non-reciprocal gene conversion in which segments of silent *pilS* loci are copied into the expressed *pilE* gene. Multiple silent *pilS* cassettes distributed throughout the chromosome provide donor sequences that replace homologous portions of *pilE*, thereby generating extensive pilin diversity while preserving expression of a single functional pilin gene. Genetic studies have established that this process requires the conserved *pilE* G-quadruplex, RecA, and numerous proteins involved in homologous recombination and DNA processing (36,37).

Although the essential role of the *pilE* quadruplex has been firmly established, its precise function during gene conversion has remained unclear. Earlier models proposed that the quadruplex promotes local DNA processing or serves as a landmark for recombination. However, the structural relationship between the quadruplex and the RecA filament has remained unknown.

The docking calculations presented above establish four observations.

1. only three-layered quadruplexes are accommodated;
2. compact loops are required;
3. the quadruplex occupies the inter-L2 region;
4. filament propagation is compatible only in one direction.

#### Proposed model

##### I. Initiation

The folded *pilE* quadruplex is incorporated into a growing RecA filament while the downstream ssDNA continues along the filament axis. The promoter-proximal region located 3′ of the quadruplex remains associated with RecA and is therefore protected during subsequent recombination (Figure 7)

**Figure 7.**
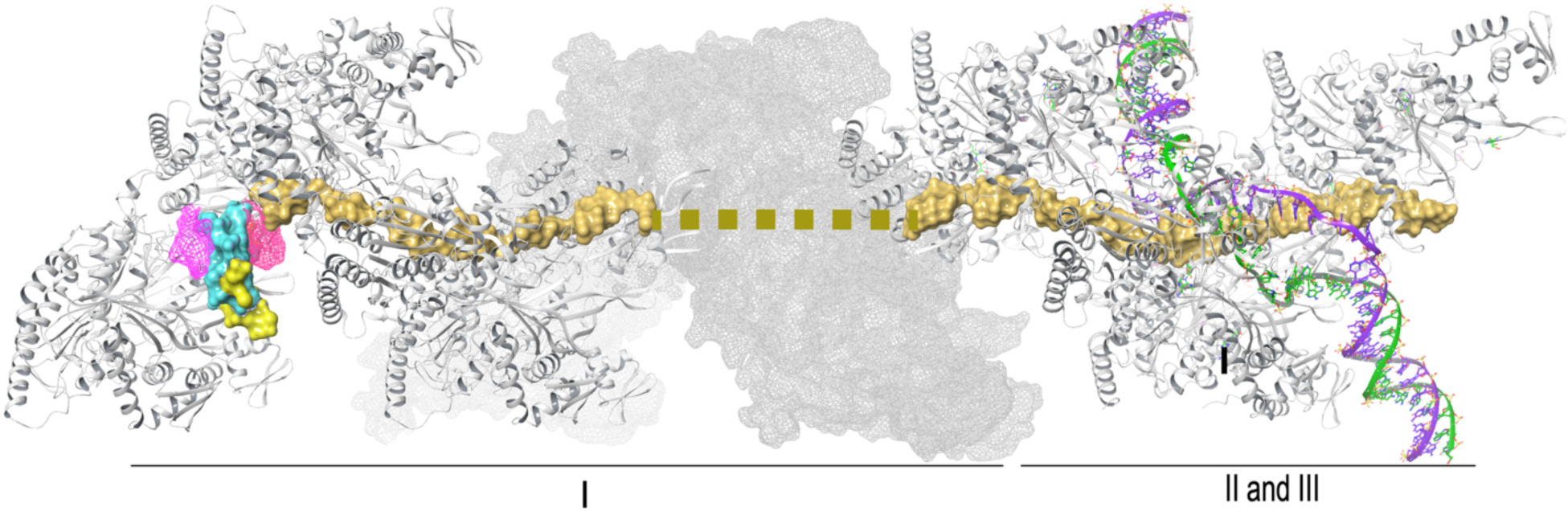
Proposed model for RecA-mediated pilE gene conversion initiated by the pilE G-quadruplex. The pilE G-quadruplex core is shown as cyan surface with propeller loops in yellow, positioned between the L2 loops of adjacent RecA monomers (magenta mesh). The RecA-bound ssDNA is shown in gold. The interrupted segment and transparent molecular surface represent continuation of the RecA filament beyond the modeled region. **I**, assembly of the RecA filament on quadruplex-containing ssDNA; **II**, homology search by the incoming pilS duplex; **III**, engagement of the homologous donor duplex and initiation of strand exchange. The pilS donor DNA is shown as purple and green ribbons. Stage I is based directly on docking into the RecA–ssDNA cryo-EM structure, whereas stages II and III represent a mechanistic model consistent with current structural and genetic data.

##### **II.** Homology search

The RecA filament performs the canonical homology search using the exposed homologous region derived from *pilE*. Consistent with recent cryo-EM studies of RecA-mediated strand exchange (46), donor duplex DNA from *pilS* engages the filament through Site II at an oblique angle before formation of a stable recombination intermediate.

##### **III.** Strand exchange

Once homology has been established, the donor duplex becomes incorporated into the RecA filament, allowing replacement of the variable *pilE* coding region while leaving the downstream promoter-associated sequence protected by the RecA filament.

##### **IV.** Resolution

Resolution of the recombination intermediate by the downstream recombination machinery restores an intact *pilE* locus containing a newly acquired donor sequence while preserving the promoter and 5′ untranslated region required for expression.

This model does not attempt to describe every enzymatic step involved in pilin antigenic variation. Rather, it provides a structural framework that integrates genetic observations, recent structural studies of RecA-mediated strand exchange, and the docking analyses presented here. The model specifically addresses how the unique geometry of the *pilE* G-quadruplex allows it to become incorporated into an active RecA filament while preserving the promoter-proximal region during gene conversion.

The structural filters identified above are not unique to *pilE*. Numerous eukaryotic recombination hotspots—including c-MYC, c-KIT, BCL2 and CEB1—also form compact three-layered parallel G-quadruplexes. Because Rad51 and Dmc1 share the RecA fold and conserve the DNA-binding architecture responsible for homologous recombination, these observations raise the possibility that the structural principles identified for the bacterial *pilE* quadruplex may represent a more general mechanism by which a restricted class of G-quadruplexes interfaces with homologous recombination.

Although genome instability at oncogenic G-quadruplex-forming loci is commonly interpreted in the context of replication-associated DNA damage and non-homologous end joining, the present structural analysis suggests that a restricted class of three-layered parallel-stranded G-quadruplexes with compact loops also possesses geometric features compatible with RecA-family recombinases. Whether analogous interactions occur with Rad51 or Dmc1 remains to be established experimentally.

## 5. Discussion

### 5.1 Structural Constraints Connect Sequence Space with DNA Architecture

The sequence-space formalism developed here identifies a highly restricted subset of DNA sequences that satisfy simultaneously the requirements of complementarity, strand symmetry, and compositional parity. Although these filtering rules are purely combinatorial, they preserve distinct classes of symmetry relationships, including mirror symmetry, reverse-complement symmetry, and shadow transformations. The present work demonstrates that these mathematical relationships are not merely abstract properties of sequence space but can correspond to physically realizable structural transitions within nucleic acids.

Among currently characterized DNA architectures, G-quadruplexes appear particularly well suited for this role because they combine structural elements that otherwise occur separately in duplex and triplex DNA.

The coexistence of parallel and antiparallel strand orientations, cis- and trans-like pairing geometries, and strand-crossing topologies enables quadruplexes to bridge symmetry relationships that appear disconnected within canonical duplex structures. In this sense, structural biology provides a physical realization of the combinatorial relationships identified in the filtered sequence space.

Importantly, the present analysis suggests that not every quadruplex is equally suited for this role. Instead, structural compatibility is governed by highly specific geometric constraints, indicating that biological selection acts not only on sequence but also on the architecture through which sequence symmetry is expressed.

An important consequence is that the filtering process does not merely eliminate sequence classes but preferentially preserves structural motifs capable of participating in higher-order DNA architectures. The present analysis suggests that G-quadruplexes represent one such surviving structural solution, linking otherwise disconnected regions of the filtered sequence space.

### 5.2 The pilE Quadruplex Represents a Structurally Specialized Recombination Intermediate

The pilE antigenic variation system provides one of the best characterized biological examples linking a G-quadruplex to homologous recombination. Our structural analysis explains several experimental observations that previously appeared independent.

First, only three-layered parallel-stranded quadruplexes are compatible with productive accommodation within the RecA filament. Second, compact propeller loops are required to position the quadruplex correctly between adjacent L2 loops. Third, modest alterations in loop size or composition produce graded reductions in compatibility that parallel experimentally observed reductions in antigenic variation. Finally, larger propeller loops or additional tetrads redirect the quadruplex toward alternative binding modes that are incompatible with the recombination-active region of the filament.

Together, these observations suggest that the native pilE quadruplex represents a highly specialized structural solution optimized for interaction with the bacterial recombination machinery rather than simply another stable G-rich secondary structure.

Although the present work focuses on the bacterial RecA system, the underlying structural principles may not be restricted to prokaryotes. The RecA family of recombinases—including Rad51 and Dmc1—shares a conserved ATPase core and homologous DNA-binding architecture despite significant evolutionary divergence. The principal structural difference relevant to the present study is the greater conformational freedom of the L2 region in the eukaryotic proteins. Preliminary docking calculations suggest that these longer loops can accommodate three-layered parallel G-quadruplexes containing somewhat larger propeller loops than are tolerated by bacterial RecA. While these observations remain preliminary, they raise the possibility that analogous quadruplex-mediated recognition mechanisms may extend beyond bacteria.

### 5.3 Geometric Selection Extends the Concept of Biological Filtering

An important outcome of the present work is the emergence of a second level of filtering beyond sequence composition.

The combinatorial framework presented above restricts the accessible sequence universe through successive mathematical filters. The structural analysis presented here indicates that an analogous process operates at the level of DNA architecture. Although numerous sequences are capable of forming G-quadruplexes, only a restricted subset possesses the geometric characteristics required for productive interaction with RecA.

Loop architecture, tetrad number, strand orientation, and overall molecular dimensions therefore constitute structural filters that further reduce the number of biologically functional conformations. The correspondence between mathematical filtering in sequence space and geometric filtering in three-dimensional structure suggests that biological evolution may exploit successive levels of constraint, progressively selecting sequences that satisfy both combinatorial and structural requirements.

From this perspective, quadruplex loop architecture constitutes an additional structural filter superimposed upon the combinatorial filters introduced earlier. Among the many theoretically possible mirror-symmetric quadruplex-forming sequences, only a small subset possesses the geometry compatible with productive interaction with the RecA filament. Functional sequence space is therefore reduced not only by nucleotide composition but also by three-dimensional architectural constraints imposed by the recombination machinery.

Taken together, the present work suggests that biologically functional DNA occupies the intersection of multiple levels of selection. Combinatorial filters determine which sequences remain accessible within sequence space, structural filters determine which of these sequences can adopt recombination-competent architectures, and molecular recognition by RecA further selects those conformations compatible with productive strand exchange.

### 5.4 Implications for Other Recombination Systems

The RecA–pilE system represents the only example currently supported by extensive genetic, biochemical, and structural evidence. Nevertheless, the geometric principles identified here may not be unique to Neisseria.

Members of the RecA recombinase family, including Rad51 and Dmc1, share conserved filament architectures and common mechanisms of homologous strand exchange. Numerous eukaryotic genomic loci associated with genome instability, including promoters of oncogenes and recombination-prone minisatellites, form predominantly three-layered parallel-stranded G-quadruplexes with compact propeller loops. Although these systems are presently interpreted primarily in the context of replication stress and double-strand-break repair, the structural compatibility demonstrated here suggests that direct interactions between selected quadruplex architectures and RecA-family recombinases deserve experimental investigation.

The present work therefore does not propose a general recombination mechanism for eukaryotic quadruplexes. Rather, it identifies a structural hypothesis that can now be tested experimentally.

### 5.5 Evolutionary Perspective

The sequence-space formalism developed here also suggests a different way of viewing homologous recombination itself. In the conventional picture, recombination primarily exchanges information between individual genomes. In a fully populated sequence space, however, recombination can equally be viewed as a mechanism that allows biological systems to navigate between neighboring regions of an otherwise fixed combinatorial landscape. Rather than creating fundamentally new sequence space, recombination selectively redistributes existing sequence combinations while preserving compatibility with structural filters.

Mirror symmetry has frequently been regarded as a simple consequence of sequence composition or local DNA symmetry. The present analysis suggests a broader interpretation.

Mirror-symmetric sequences survive successive combinatorial filters and simultaneously possess the capacity to generate higher-order DNA architectures capable of mediating genome rearrangement. Rather than representing isolated structural curiosities, these motifs may constitute evolutionary solutions that couple information storage with controlled structural plasticity.

In this framework, recombination is not viewed solely as a biochemical process acting upon DNA sequence but also as a phenomenon constrained by the geometry of the structures that particular sequences are able to adopt.

The present work integrates three levels of description that are rarely considered together: combinatorial organization of sequence space, structural properties of G-quadruplex DNA, and molecular mechanisms of homologous recombination. While each component has previously been studied independently, their convergence suggests that certain secondary structures occupy privileged positions within sequence space because they satisfy multiple structural constraints simultaneously. The pilE system provides one experimentally validated example. Whether similar principles contribute to recombination at eukaryotic G-quadruplexes remains an important question for future investigation.

Mutation is commonly viewed as the principal mechanism by which biological systems explore sequence space. The present framework suggests that homologous recombination may instead be interpreted as a mechanism for navigating an already existing combinatorial landscape. This perspective may be particularly relevant to horizontal gene transfer in bacteria, where mobile genetic elements continually introduce pre-existing sequence variants into recipient genomes. Successful incorporation of these variants depends not only on sequence homology but also on compatibility with the structural constraints imposed by the recombination machinery. In this view, recombination functions less as a generator of sequence diversity than as a navigator through the accessible regions of sequence space.

### 5.6 Concluding Remarks

The principal contribution of this work is the integration of combinatorial sequence analysis with structural biology. Beginning from a mathematically filtered representation of sequence space, we identify classes of symmetry relationships that are subsequently realized by specific DNA architectures and finally connected to experimentally characterized biological systems.

This progression—from sequence symmetry, through molecular structure, to homologous recombination—suggests that the organization of genomic sequence space reflects not only statistical constraints but also structural possibilities available to DNA itself. The pilE G-quadruplex illustrates one experimentally supported realization of this principle. Whether analogous structural solutions have evolved elsewhere in biology remains an open question, but the geometric framework presented here provides a basis for addressing that question experimentally.

Viewed from this perspective, sequence space is not merely a combinatorial universe populated by DNA sequences. It is a structural landscape whose accessible trajectories are progressively constrained by molecular architecture, and whose biologically meaningful pathways emerge only where sequence symmetry, DNA structure, and recombination machinery converge.

