## Supplementary Material for "Recombinogenic G-quadruplexes in the Newtonian DNA Sequence Space"

Vitaly Kuryavyi

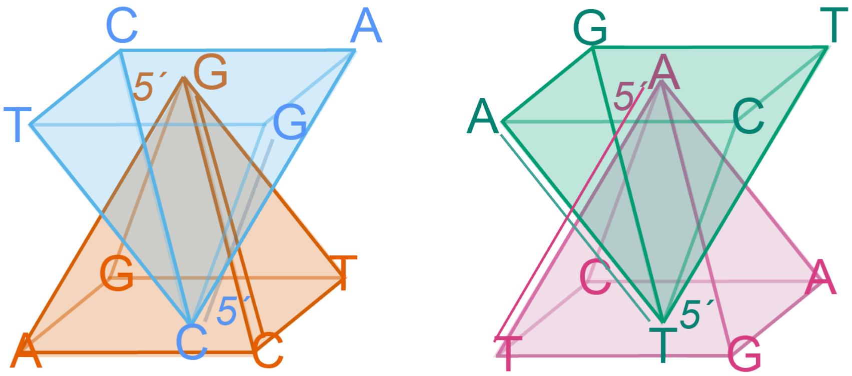


**Figure S0. Pairing of dinucleotide pyramids according to complementary symmetry.** Dinucleotide pyramids are grouped into complementary GC and AT pairs, with one pyramid in each pair inverted relative to the other. The edges corresponding to 5′-GC-3′ and 5′-CG-3′ (left panel), and to 5′-AT-3′ and 5′-TA-3′ (right panel), are shown as double lines to indicate the degeneracy of complementary interactions that arises for specific sequence classes of even length. This degeneracy underlies the occurrence of complementary sequence pairs occupying identical positions in the corresponding sequence-space pyramids.

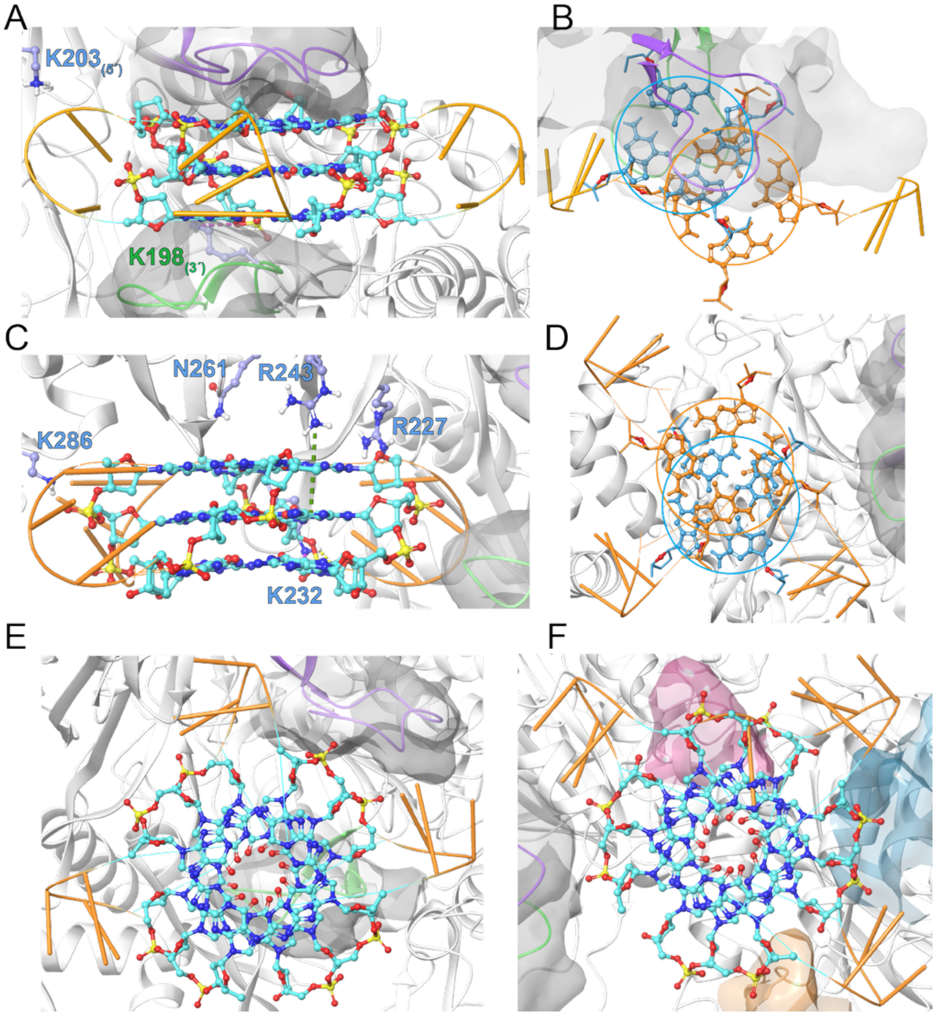

**Figure S1. Docking poses of a three-layered parallel-stranded G-quadruplex with long propeller loops (human telomeric topology). (A,B)** An inverted docking pose positioned between the L2 loops of adjacent RecA monomers. **(A)** Side view showing displacement of the quadruplex core away from the helical axis of the RecA filament. **(B)** Top view of the same complex. The telomeric quadruplex is shown in orange; a G-tetrad from the docked pilE quadruplex (blue) is superimposed for comparison, illustrating the lateral displacement imposed by the enlarged propeller loops. **(C,D)** Docking pose within the cavity formed by the C-terminal domain (CTD) and the L2 region. **(C)** Side view. **(D)** Top view. Relative to the loop-free four-layered quadruplexes (one reference tetrad shown in blue), the bulky propeller loops shift the quadruplex toward the CTD, preventing productive insertion between the L2 loops. **(E)** Docking pose oriented approximately perpendicular to the L2 interface. **(F)** Alternative pose shifted toward the L1 region. In all panels, the L2 loops of the adjacent RecA monomers are shown in lilac and green to illustrate the topography of the recombination-active region. None of the observed docking modes reproduces the productive inter-L2 binding geometry characteristic of the native pilE G-quadruplex.

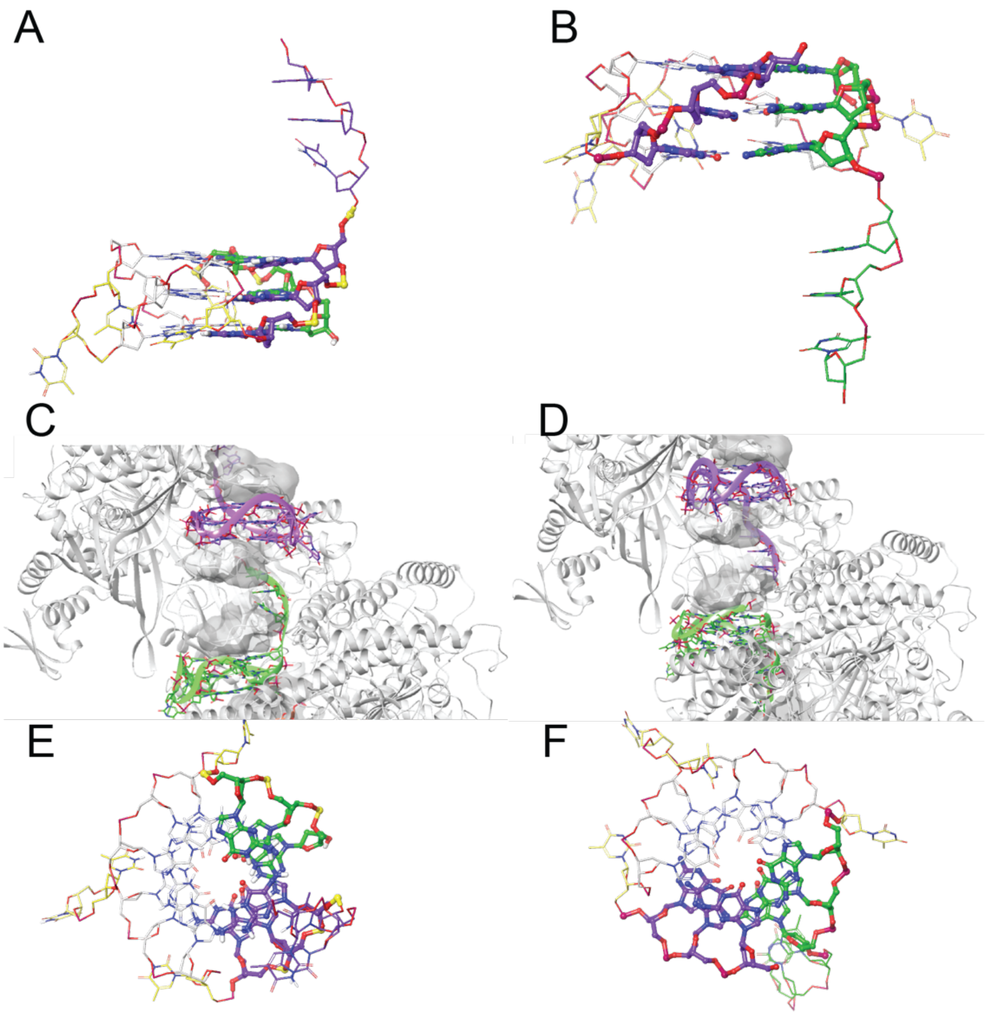


**Figure S2. Structural consequences of extending the pilE G-quadruplex within the RecA filament. (A,B)** The pilE G-quadruplex extended by single-stranded DNA at the **5′ end** (**A**) or **3′ end** (**B**). The 5′ and 3′ terminal strands of the quadruplex are colored lilac and green, respectively, matching the color scheme used throughout the manuscript. **(C,D)** Docked 5′- and 3′-extended quadruplexes positioned in adjacent inter-L2 binding sites of the RecA filament. The two complexes are separated by an inter-L2 region occupied by the intervening RecA-bound ssDNA, illustrating that successive quadruplex-containing binding sites can be connected through the regular trinucleotide organization of the filament. **(E,F)** Top view of the two extended quadruplexes demonstrates that the 5′- and 3′-extended conformations differ by an approximately one-tetrad rotational register of the quadruplex core. This rotational relationship provides the structural basis for the alternative orientations of the quadruplex observed within the RecA filament.

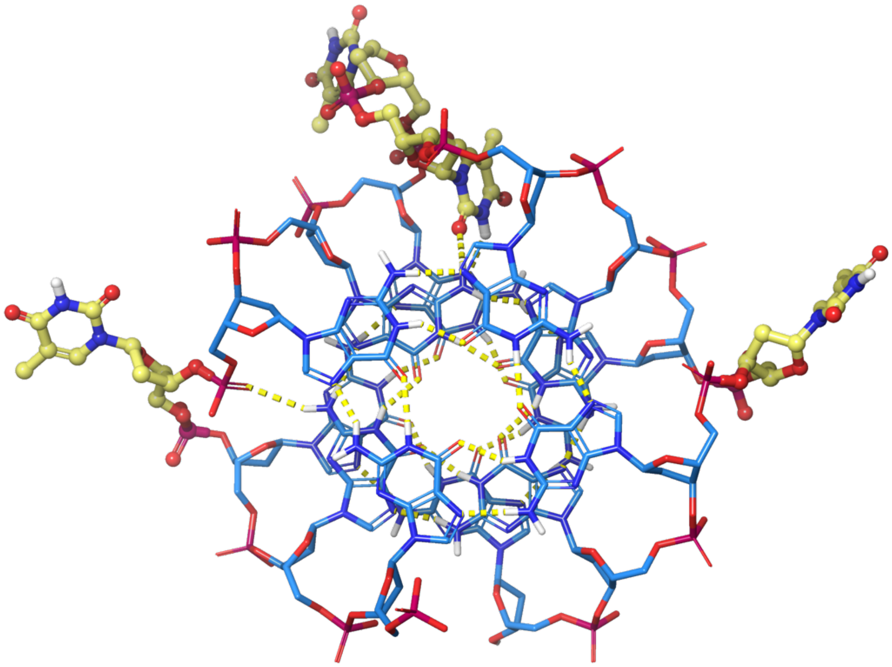

**Figure S3. Top view of the monomeric pilE G-quadruplex.** The top projection emphasizes the compact architecture of the pilE G-quadruplex and separates the molecule into four structural sectors corresponding to the three propeller loops and the 5′–3′ groove. The two single-residue loops (dT4 and dT13) and the compact two-residue dT8–dT9 loop occupy three sectors of the molecule, whereas the remaining sector corresponds to the uninterrupted 5′–3′ groove. This arrangement produces a highly asymmetric molecular surface that is preserved in the RecA-bound docking models and contributes to the reproducible orientation of the pilE G-quadruplex within the inter-L2 binding site.
